# Spatial transcriptomics unveils immune cellular ecosystems associated with patient survival in diffuse large B-cell lymphoma

**DOI:** 10.1101/2024.09.16.613252

**Authors:** Alba Díaz Herrero, Hector Fernando Pelaez-Prestel, Lucile Massenet-Regad, Maëva Veyssiere, Julien Calvani, Caterina Cristinelli, Jacqueline Lehmann-Che, Véronique Meignin, Catherine Thieblemont, Véronique Blanc, Vassili Soumelis, Pierre Tonnerre

**Affiliations:** Institut de Recherche Saint-Louis, Université Paris-Cité, Inserm U976 HIPI, F-75010 Paris, France; School of Medicine, Department of Immunology, Complutense University of Madrid, Madrid, Spain; Pathology department, hôpital Saint-Louis, AP-HP, F-75010 Paris, France; Université Paris Cité, 85 boulevard St Germain, F-75006 Paris, France; Assistance Publique – Hôpitaux de Paris, Hôpital Saint-Louis, Hemato-oncologie, 1 Avenue Claude Vellefaux, F-75010 Paris, France; Inserm U1153, Hôpital Saint-Louis, 1 avenue Claude Vellefaux, F-75010 Paris, France; Institut de Recherches Servier, Gif-sur-Yvette, France; Department of Immunology-Histocompatibility, Saint-Louis Hospital, AP-HP Nord, Université Paris Cité, F-75010 Paris, France

## Abstract

Diffuse Large B-cell Lymphoma (DLBCL) is the most prevalent subtype of non-Hodgkin’s lymphoma for which current therapeutic strategies remain insufficient. The diffuse nature of DLBCL, lacking distinct tissue structures, represents a challenge to elucidate the cellular organization and interactions within the tumor microenvironment (TME). In this study, we applied spatial transcriptomics to identify spatially-resolved gene expression profiles in 10 DLBCL tissue samples, identifying distinct immune cell infiltration and colocalization patterns. These profiles were classified into six cellular ecosystems (Cell-Eco) that differ in cellular composition, functional patterns, and neighborhood characteristics. The spatially-resolved Cell-Eco signatures provided prognostic scores that stratified patients with different overall survival rates. We also found that C1q+ tumor-associated macrophages are the primary cells interacting with malignant B cells and influencing the spatial architecture of the TME. This study provides novel biological insights into the complexity of the TME in DLBCL and highlights the potential prognostic value of its spatial organization.

**Graphical abstract:** 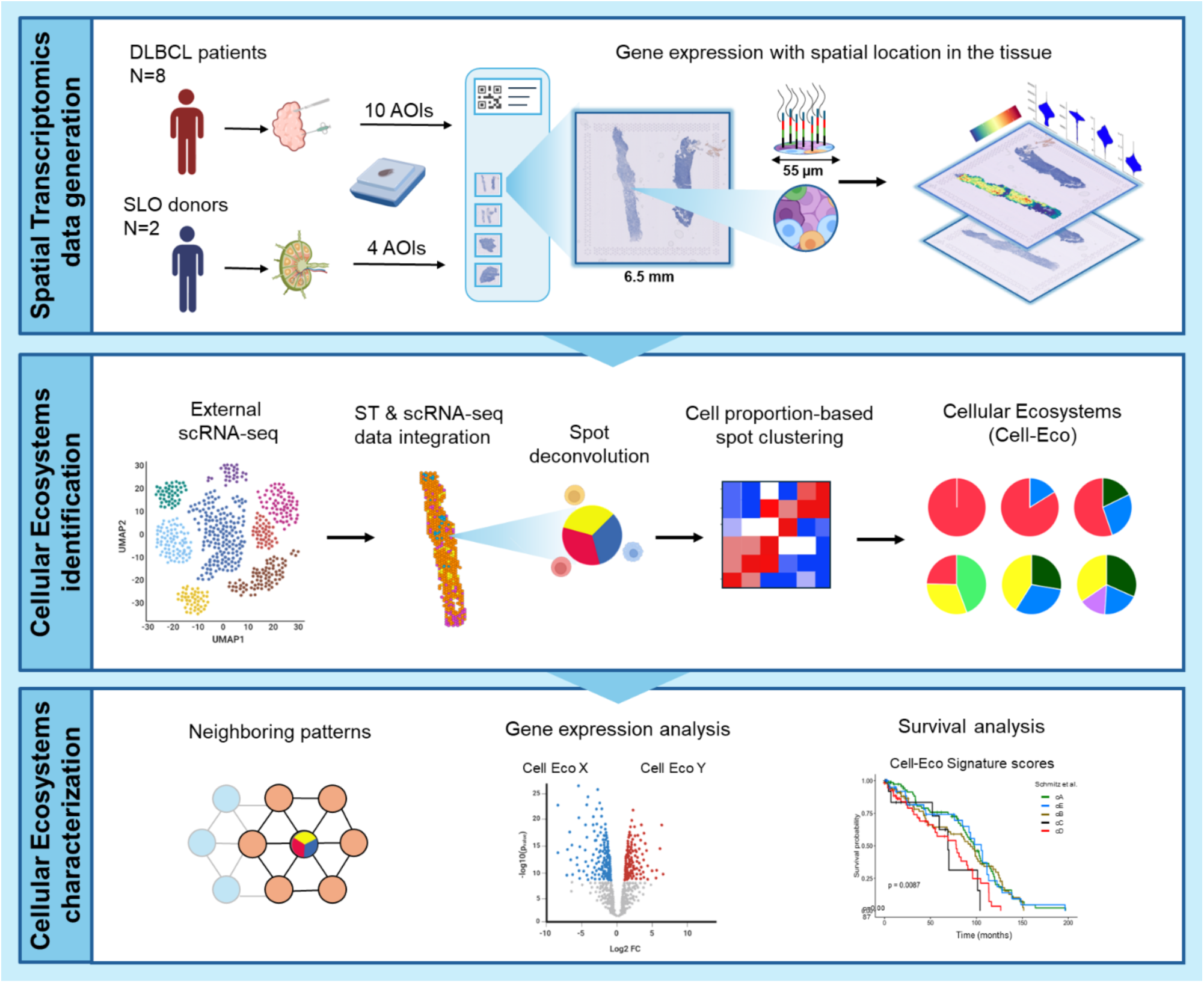

**Key findings:**

- Spatial transcriptomics classifies DLBCL tissues based on immune cell infiltration and colocalization patterns.
- DLBCL tumor microenvironment consists of cellular ecosystems (Cell-Eco) that differ in cellular composition, transcriptomic profiles and neighborhood characteristics.
- Spatially-resolved Cell-Eco signatures stratify patients with different overall survival.
- C1q+ tumor-associated macrophages primarily interact with malignant B cells and contribute to the spatial organization of the tumor microenvironment.

## Introduction

Diffuse large B-cell lymphoma (DLBCL) is the most common subtype of Non-Hodgkin’s Lymphoma (NHL) worldwide and accounts for about 40% of all cases^1,2^. Despite the significant improvements of patient outcomes with the first line treatment R-CHOP (rituximab, cyclophosphamide, doxorubicin prednisone and vincristine), approximately 40% of the patients experience relapse or become refractory^3^. A better understanding of the biological mechanisms associated with disease progression is essential to develop new therapies with increased response rate and improved overall survival.

The study of the tumor microenvironment (TME) has become of great importance in elucidating the factors that contribute to disease progression and relapse in DLBCL. Both the cellular composition and spatial organization have been linked to disease stage and response to treatment in various solid cancers^4–6^. However, a hallmark of DLCBL is its diffuse pattern, characterized by the scattered distribution of malignant cells within the TME^2,7,8^, hindering a comprehensive assessment of the spatial architecture and associated cell states in DLBCL. Previous studies investigating the cellular composition and gene expression profiles of the TME in DLBCL have provided important insights into the heterogeneity of immune cell infiltration and its connection to tumor progression^9–11^, treatment resistance and clinical outcomes^9,12–15^,. Nevertheless, these characterizations were limited by the lack of spatial context of the immune cell profiles, including their localization within the TME and the associated communication pathways. Recent studies have integrated tissue coordinates into detailed analysis of the DLBCL TME, using high parameter imaging^16–18^ or digital spatial profiling of macrophages^19^. However, these studies faced technical constrains such as the limited number of parameters analyzed and a narrow focus on specific immune cell types. A comprehensive characterization of the different cellular immune profiles within the tissue topology, integrating spatial localization, cellular interactions and communication patterns, is still missing in DLBCL.

In this study, we used spatial transcriptomics to generate a precise mapping of gene expression within the tissue architecture of DLBCL. We show that DLBCL tissues are structured into cellular ecosystems (Cell Eco) with distinct immune cell compositions, functions, and communication networks. In addition, we identify Ecosystem-associated transcriptomic signatures that are linked to patient overall survival. This study highlights the importance of dissecting the TME spatial immune organization for the identification of clinically relevant targets or biomarkers.

## Results

### Spatially-resolved transcriptomic profiles categorize DLBCL tissues by immune cell infiltration and co-localization patterns

We profiled gene expression with morphological context in a total of 10 DLBCL and 4 secondary lymphoid organs (SLO) tissue sections from formalin-fixed, paraffin-embedded (FFPE) blocks, using Visium Spatial Gene Expression technology from 10x Genomics (Figure 1A). This technology allowed the spatial mapping of tissues as 55μm spots with associated transcriptomic information.

**Figure 1:**
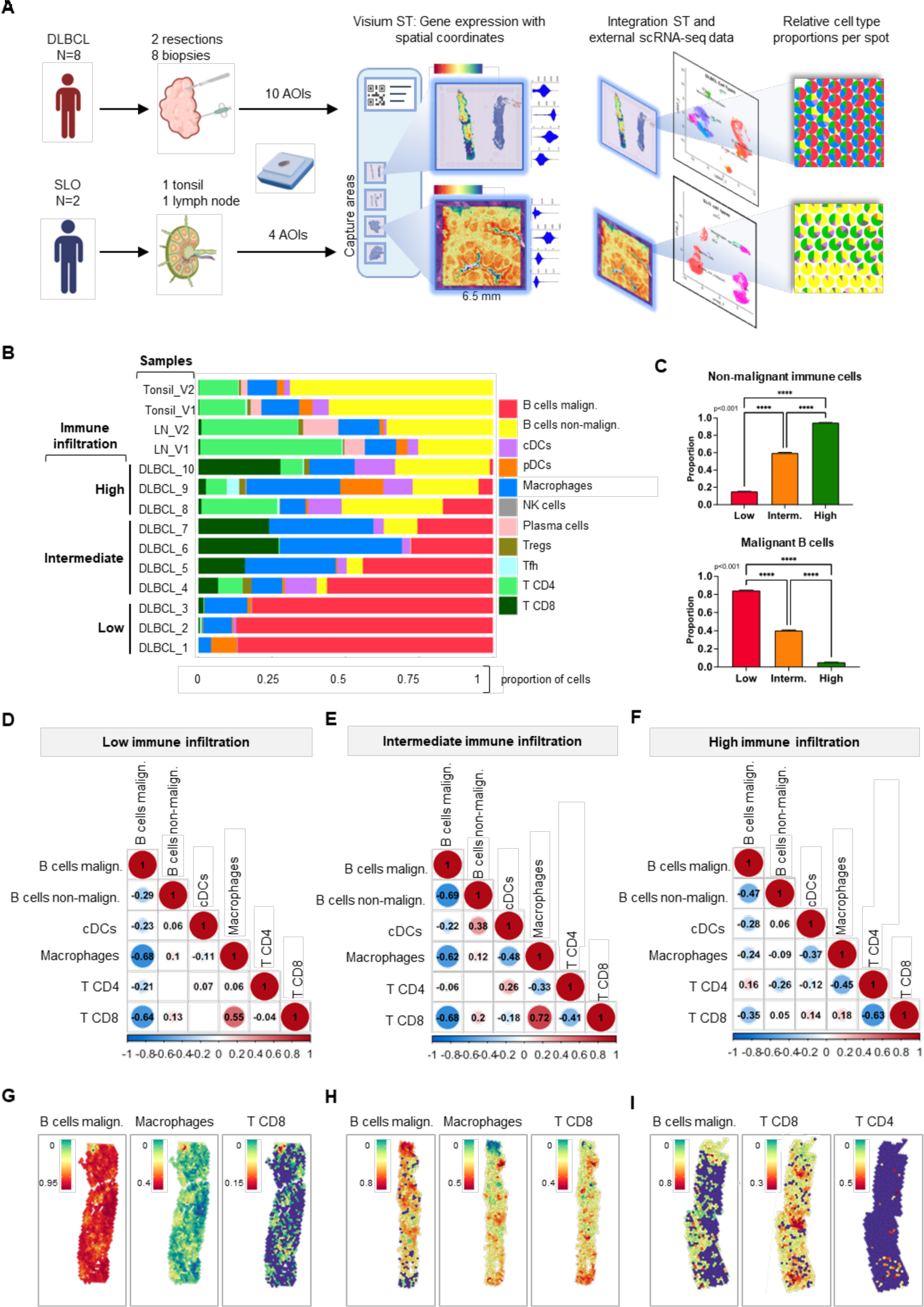
Spatially resolved transcriptomic profiles classify DLBCL tissues by immune cell infiltration and co-localization patterns. A) A schematic workflow of ST data generation with Visium (10X Genomics). All the samples were formalin-fixed and embedded in paraffin. After RNA quality control, samples were placed in Visium Spatial slides, which include 4 capture areas of 6.5 × 6.5 mm2, with 5000 barcoded spots. ST sequencing results is a matrix of gene counts and coordinates, which translate into gene expression profiling with morphological context. Single Cell Reference data from publicly available data sets is annotated and integrated with ST data to perform spot deconvolution, providing the relative cell type proportions per spot. B) Bar plot representing the mean proportions of RCTD-deconvoluted immune cell-types per ST sample. DLBCL ST samples are divided into 3 groups based on their non-malignant immune infiltration. Statistical differences of cell-types proportions across samples are represented in Figure S3D. C) Mean comparison of malignant B cells proportions (above) and non-malignant immune cells (bellow) per DLBCL sample group. The immune infiltrate compartment is represented as the sum of mean relative cell-type proportions other than malignant B cells from the RCTD deconvolution outputs. Multiple comparisons were performed using Kruskal-Wallis statistical test and Dunn’s post-hoc (∗∗∗∗p < 0.0001). D-F) Intra spot cell-type colocalization patterns per DLBCL sample group. Color intensity and circle size represent the Pearson correlation coefficients: red indicates positive correlations and blue indicates negative correlations. Only significant correlations (p<0.05) are shown. G-I) Deconvoluted proportions of general immune cell types in one representative DLBCL ST sample of the Low (G), Intermediate (H) and High (I) immune infiltrated groups.

DLBCL samples were collected from eight different patients, including two resections and eight biopsies (n=10 in total); n=6 pre-treatment biopsies from six different patients, n=2 longitudinal biopsies (before and after R-CHOP therapy) from an additional patient, and n=2 different pre-treatment biopsies from another patient. Samples were obtained from the Pathology Department at Hospital Saint Louis in Paris, France (Table S1). Additionally, two adjacent sections from a healthy lymph node (LN) and two from a non-metastatic tonsil were collected from individuals undergoing non-cancer-related surgeries at Hospital Bichat and Robert Debré in Paris, France.

First, to profile intra-spot immune cell composition, we integrated public single-cell RNA sequencing data (scRNA-seq)^14,15^ (Table S2), with our in-house generated spatial transcriptomics (ST) data to deconvolute cell-type mixtures present in each spot (Figure 1A). scRNA-seq data from the CD45+ compartment of DLBCL and SLO tissues were manually curated and annotated based on well-established markers in the literature (Figure S1A, S1B, S1C, S1D and Table S3). Annotation of malignant B-cells was based on irregular copy number variation. Subsequently, this scRNA-seq reference data was integrated with the corresponding ST DLBCL or SLO sample group to ensure sample type matching. Cell type deconvolution was performed using robust cell-type decomposition (RCTD)^20^, one of the best performing computational methods for spatial transcriptomics integration-based deconvolution ^21,22^. As a result of this integration, cell type proportions were assigned to each individual spot (Figure 1A, Figure S1E and S1F; Figure S2A).

The robustness of RCTD decomposition was assessed on control tissue. This was done by cross-comparing the deconvolution outputs for each region (Figure S1F), the spot-cluster annotation based on differentially expressed genes (DEGs) (Figure S1G and S1H) and the histological annotation from SLO. The histological annotation was revised by expert pathologists (Figure S1I). Both DEGs per cluster and decomposed cell subsets proportions were aligned with the histological annotation, as well as with the gene expression profiles and cell composition previously described for each anatomical region^23–25^. This consistency confirmed the reliability of RCTD cell-type deconvolution for spots in spatial transcriptomics samples.

To validate the tissue topology at the protein level, we conducted immunofluorescence staining for selected markers that represent major immune cell types. This was done on adjacent sections from the same FFPE blocks used for spatial transcriptomics (Figure S2B). We mapped the expression of three lineage-associated markers (CD20, CD3 and CD68) at the protein level and compared it to the decomposed cell type identification from spatial transcriptomic data of each sample. We observed similar localization patterns between RCTD-decomposed cell types and the fluorescence signals of their corresponding markers in the tissue structure (Figure S2A and S2B). Furthermore, we found correlation between the frequency of cells identified by RCTD decomposition and the frequency of cells expressing the corresponding markers at the protein level (Figure S2C).

Subsequently, we investigated the spot-based proportions of deconvoluted immune cells across control and DLBCL samples. Eleven populations of immune cells were identified, including malignant B cells, non-malignant B cells, conventional dendritic cells (cDCs), plasmacytoid dendritic cells (pDCs), Macrophages, natural killer cells (NK cells), plasma cells, regulatory T cells (Tregs), follicular T helpers (Tfh), CD4+ T cells (T CD4), CD8+ T cells (T CD8). Inferred proportions of immune cells per spot varied considerably across individual samples (Figure 1B and Figure S2D), confirming the heterogeneity of DLBCL tissues based on immune cell infiltrates. Based on the immune infiltration proportion tertiles (tertile 1=0.4362144, tertile 2= 0.7433377), we classified the DLBCL samples into three groups: low, intermediate and high immune cell infiltration. In these groups, the proportions of immune infiltration and malignant B cells were inversely related (Figure 1C).

Then, we investigated the colocalization patterns among sample groups by conducting a correlation analysis on the frequencies of deconvoluted cell types within each spot. We observed that for the three groups, malignant B cells showed negative or no correlation with any other immune cell types, indicating potential mechanisms of immune evasion (Figure 1D, 1E and 1F). For low and intermediate immune cell infiltrated samples, CD8+ T cells and macrophages displayed a significant spatial correlation together, suggesting co-localization patterns. Interestingly, the presence of these cells was anticorrelated with malignant B cells, suggesting immune cell niches within the tumor microenvironment (Figure 1D, 1E, 1G and 1H). CD4+ T cells presence is more prominent in the high immune-infiltrated group and anticorrelated to CD8+ T cells and macrophages, highlighting different co-localization patterns and cellular co-occurrence arrangements in the tissue (Figure 1F and 1I).

### Immune cells are distributed into cellular ecosystems with different neighborhood characteristics

To characterize spatial identities based on colocalization patterns, we used the Leiden algorithm, an improved version of Louvain unsupervised clustering ^26^. It allowed the grouping of a total of 7824 spots with similar immune cell-type composition across ten DLBCL samples (Figure 2A). We identified six different clusters defining distinct cellular ecosystems (Cell-Eco). These Cell-Eco are characterized by distinct cellular compositions and abundances across the tissue samples (Figure 2A and 2B; Figure S3A, Table S4). Among these Cell-Eco, three were characterized by a predominance of malignant B cells, with or without the presence of macrophages and CD8+ lymphocytes. As expected, Cell-Eco with a higher proportion of malignant B cells (Cell-Eco 1 and 2) were more frequent among the low immune infiltration samples (Figure S3A and S3B). Cell-Eco 3 showed its highest abundance in individuals with intermediate immune infiltration. (Figure S3B). In contrast, Cell-Eco 4, 5 and 6 were present at higher frequencies in highly immune infiltrated samples (Figure S3B), although they were different in immune cell compositions. Notably, Cell-Eco 4 was the only Cell-Eco with significant presence of CD4+ T cells, together with both malignant and non-malignant B cells, which could be reminiscent of the healthy lymph node cellular composition. Cell-Eco 5 and 6 included non-malignant B cells together with CD8+ T cells and macrophages, and only differed by the presence of cDCs in Cell-Eco 6 (Figure 2B; Figure S3A). The cellular ecosystems define common identities among spots, which represent spatial units of cells that share coordinates in the ST gene count matrix. However, they also exhibit specific distributions across the tissue section. We studied the compartmentalization of samples groups by analyzing their Cell-Eco neighboring patterns. For that purpose, we calculated a neighbor score based on the relative frequency of Cell-Eco in direct vicinity by pairs, and then determined the mean neighbor scores per sample group (Figure 2C-E). We observed that in low infiltrated samples, malignant B cell-rich spots (Cell-Eco 1 and Cell-Eco 2) form aggregates not only at a spot level (Figure 1D-F) but also at a broader scale as indicated by their higher neighbor score (Figure 2C). The high neighbor score of Cell-Eco 3 for the intermediate infiltration group of samples can be attributed to their group-specificity and abundance, which likely contributes to their spatial proximity (Figure 2B and 2D; Figure S3B). Cell-Eco 4 displays great aggregation scores (0.16) in the high infiltrated group, which could also refer to their abundances (Figure 2E). Notably, Cell-Eco 5 and 6 are the ecosystems in both intermediate and high immune infiltration groups that showed a high dispersion pattern, as indicated by their low Mean neighbor scores among different pairs (Figure 2D and 2E). Overall, these data indicate that the different Cell-Eco exhibit distinct neighboring patterns, revealing compartmentalization based on cellular composition in DLBCL tissue samples.

**Figure 2:**
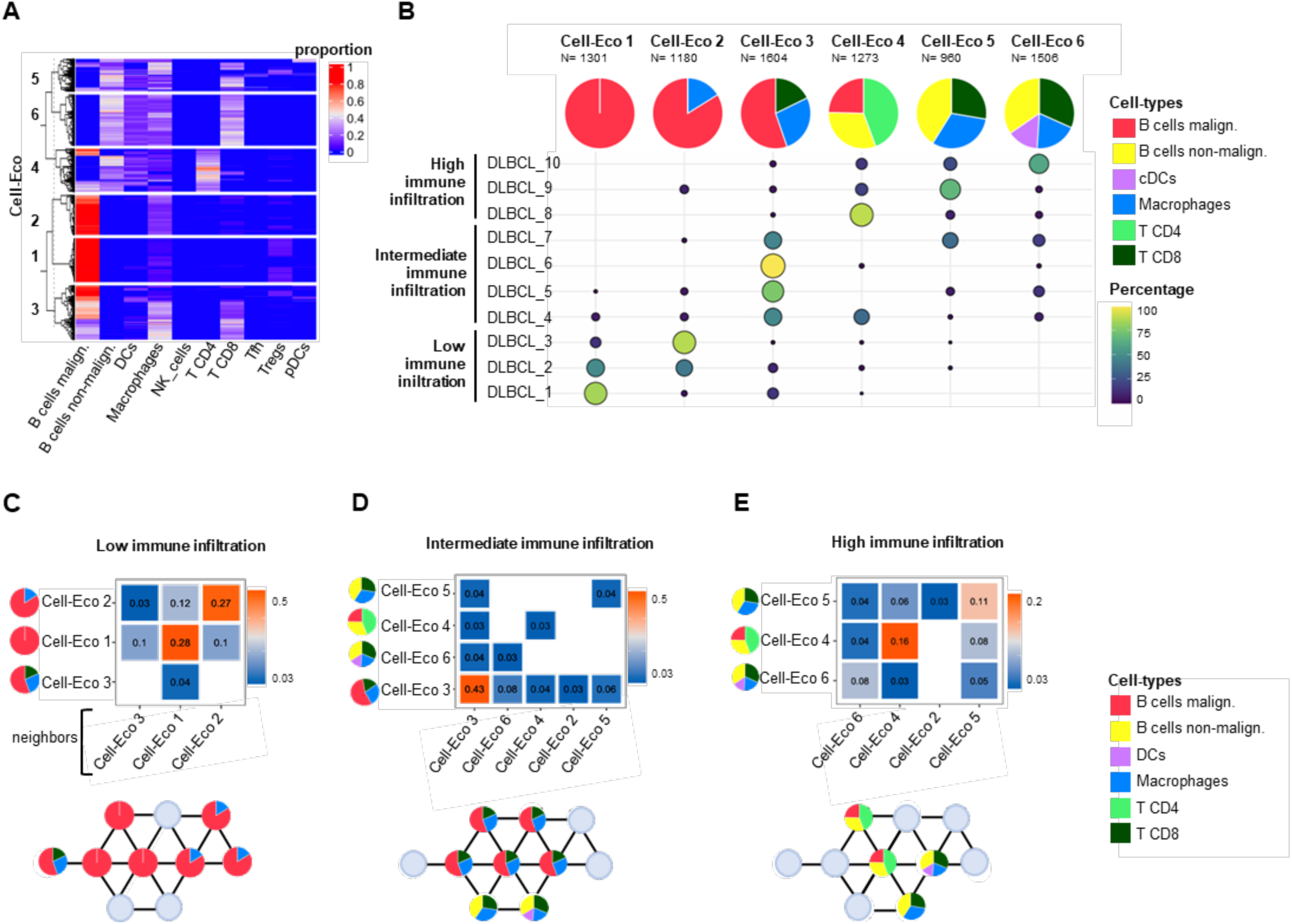
Cellular Ecosystems and neighborhood characteristics in DLBCL tumor microenvironment. A) Heatmap of deconvoluted cell-type proportions per spot of 10 ST DLBCL samples, clustered by the by Leiden algorithm (resolution 0.2). B) Above: Pie charts representing mean cell-type proportions of each Leiden cluster (Cell-Eco). Proportions lower than 10% were excluded from the analysis and means were scaled to sum 100% per Cell-Eco. Bellow: bubble plot indicating the percentage of each Cell-Eco per sample. C-E) Heatmaps of mean neighbor scores (above) of Cell-Eco pairs with their schematic representation (below) of samples of the Low (C), Intermediate (D) and High (E) immune infiltrated groups. The heatmap represents mean neighbor score values; high in orange and low in blue. Only scores higher than 0.03 are represented. Equations describing the score calculations are detailed in the methods. Schemas represent Cell-Eco neighboring frequencies based on the scores, in a virtual area of 10 spots. Gray dots indicate negligible neighbor presence.

Interestingly, different biopsies (DLBCL_5 and DLBCL_7) from the same patient (p04), both collected from the same procedure and prior R-CHOP treatment, consistently presented high frequencies of Cell-Eco 3 and fell into the same Intermediate group, indicating uniform cellular composition across the two different biopsies (Figure 1B, Figure S3C, Table S4). Conversely, biopsies DLBCL_2 and DLBCL_3 from patient p02, collected 232 days apart before and after R-CHOP treatment respectively, exhibited different Cell-Eco compositions. More specifically, DLBCL_3 collected at relapse, showed a higher frequency of Cell-Eco 2 (86.24%) compared to the tissue sample collected at baseline (41.08%) (Figure 1B, Figure S3C, Table S4), suggesting a possible remodeling of the cellular ecosystems after treatment.

### Cellular ecosystem signatures allow the stratification of DLBCL patients with different overall survival scores

To evaluate whether spatially resolved Cell-Eco signatures hold a potential prognostic value for DLBCL patient outcomes, we performed survival analysis on two publicly available bulk RNA sequencing (RNA-seq) datasets. The dataset from Schmitz et al.^27^ and Visco et al.^28^ included RNA-seq data from biopsy samples collected at diagnosis in n=277 patients, and n=241 respectively, who had survival information over a period of 200 months.

We scored Cell-Eco gene signatures from the ST data, based on differential expression analysis (Table S5) and clustered the RNA-seq data patients accordingly. Remarkably, Cell-Eco signature scores allowed the stratification of the patients from Schmitz et al.^27^ cohort into five distinct clusters (cA-cE) with different overall survivals (OS) Figure 3A and 3B). Patient groups cA and cE were associated with better OS (Figure 3B) and displayed high scores of Cell-Eco 5, in combination with other non-malignant Cell-Eco signatures (Cell-Eco 4 for cA and Cell-Eco 6 for cE subgroups) (Figure 3C). Interestingly, high score of Cell-Eco 5 was also associated with greater OS for patients from Visco et al.^28^ (Figure S4A and S4B). On the other hand, patients with worse OS in the Schmitz et al.^27^ cohort, belonging to the clusters cC and cD (Figure 3B), exhibited high scores of Cell-Eco 3 and 2, or Cell-Eco 3 and 1, respectively (Figure 3C).

**Figure 3:**
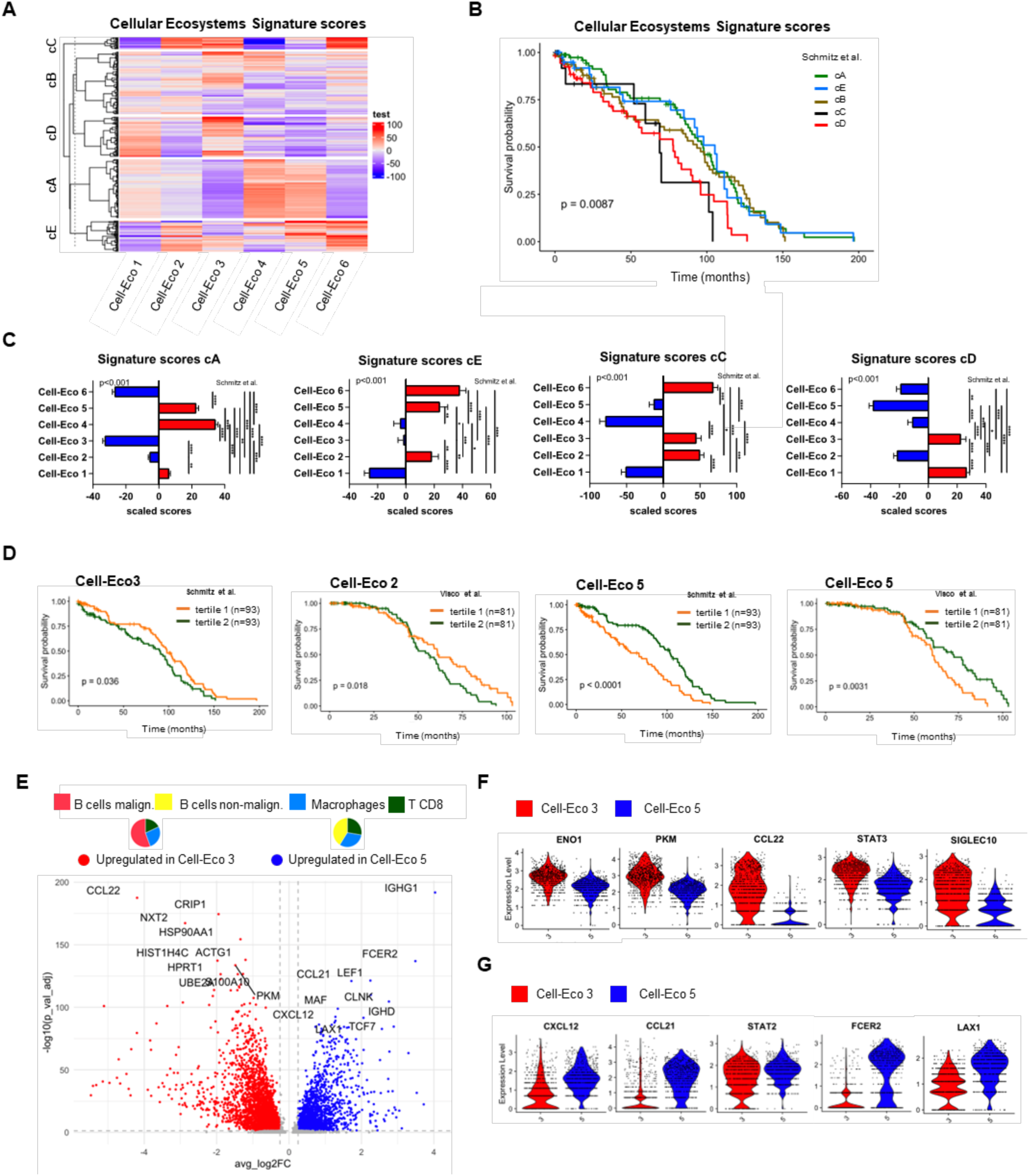
Cellular Ecosystems signatures allow stratification of patients with different overall survival scores. A) Heatmap of patient stratification based on Cellular Ecosystems signature scores in the Schmitz cohort of DLBCL RNAseq data, revealing five distinct clusters (cA=80, cB=85, cC=16, cD=54, cE=42). B) Kaplan-Meier curves showing the overall survival of the patient clusters from Figure 4A showing the p value from Log-rank test. C) Bar plots of Cell-Eco signature scores for individual patient clusters (cA, cE, cC, cD). D) Kaplan-Meier curves showing the overall survival of patients based on individual Cell-Eco signature scores in two individual RNAseq datasets. P-values generated by Log-rank test. E) Volcano plots showing differentially expressed genes between Cell-Eco 3 (in red) and in Cell-Eco 5 (in blue), based on log2FC>0.25 and adjusted p-value<0.05. P-values were determined by the Wilcoxon-rank sum tests. Labels of the top 10 genes with lower adjusted p-values are displayed in the plot. F-G) Violin plots showing genes differentially upregulated in Cell-Eco 3 (F) and Cell-Eco 5 (G).

We also examined the influence of Cell-Eco individual signatures on OS. Survival analysis was performed on patients grouped by tertiles of individual Cell-Eco signature scores. We observed high signature scores of Cell-Eco 3 and 2 in patients with lower OS in Schmitz et al.^27^ and Visco et al.^28^ cohorts, respectively. Importantly, Cell-Eco 5 was associated with better survival for DLBCL patients in the two independent cohorts (Figure 3D).

Despite the opposite impact on survival, Cell-Eco 3 and 5 present similar cellular compositions that differ only in the malignancy status of the B cells (malignant B cells for Cell-Eco 3 and non-malignant B cells for Cell-Eco 5). Differential gene expression analysis was conducted to explore further differences between these two Cell-Eco at the transcriptomic level. This analysis revealed a strong glycolytic profile in malignant Cell-Eco 3. This profile has been previously linked to progression and resistance, and is upregulated in DLBCL malignant B cells (PKM, STM1, ENO1 and CDK1)^29^. This Cell-Eco also exhibits higher expression of cell cycle, proliferation (HIST1H4C, NXT2, CRIP1) and cell motility genes (ACTG1, ACTB)^30^ compared to Cell-Eco 5 (Figure 3E and 3F; Table S6). In addition, we noted a significant upregulation of CCL22 in Cell-Eco 3, a chemokine produced by tumor-associated macrophages which has been linked to tumor metastasis and reduced patient survival^31^. Other genes highly expressed in Cell-Eco 3 included genes associated with anti-inflammatory signaling (STAT3, SIGLEC10, CD24)^32,33^. Conversely, Cell-Eco 5 associated with higher expression of immune cell recruitment genes (CXCL12, CCL21 and CCL19)^34^ and IFN-induced transcription factors (STAT2, STAT4), together with genes associated with macrophage functions (FCER2, MAF) as well as T cell differentiation and functions (LAX1, TCF7, CD40LG, ILR7)^35,36,37^ (Figure 3E and 3G; Figure S4C Table S6). Altogether, these findings underscore the importance of assessing the biological mechanisms involved in each cellular ecosystem, beyond their immune cell composition.

### Tumor-supportive macrophages are the predominant cell type in direct contact with malignant B cells and shape the DLBCL tumor microenvironment spatial architecture

To further investigate the role of the macrophages located in highly malignant microenvironments, we performed a differential gene expression analysis (DEA) between Cell-Eco 1 and Cell-Eco 2, both composed mostly of malignant B cells but differing by the presence of macrophages in Cell-Eco 2. (Figure S3B). We observed that 2332 and 4473 genes were significantly upregulated (adjusted p-value<0.05 and log2FC>0.25) in Cell-Eco 1 (B cells malignant mean _Cell-Eco 1_= 100%,) and Cell-Eco2 (B cells malignant mean _Cell-Eco 2_= 83.68%, macrophages mean _Cell-Eco 2_=56.82%), respectively. The high number of DEGs underscores the differences in gene expression profiles between the two entities (Table S7). Cell-Eco 1 presents a transcriptomic profile that reflects the predominant composition of malignant B cells. This phenotype is supported by the DisGeNET enrichment analysis, a data base of gene-disease association, which identified the top 50 significant genes of the DEA as an Adult Diffuse Large B-Cell lymphoma signature (Figure S5A). Cell-Eco 1 also presents significantly higher expression of genes associated to B cell memory and plasma cells (TNFRSF13B, IGHM, IGHG3, CCR7, IL10RA, IRF4, PRDM1, JAK1) which indicates a post-GC differentiation status^38,39^. Conversely, some marker expressions of germinal center dark zone like MS4A1, AIDA and FOXP1^38^, are increased in Cell-Eco1. It also exhibits upregulation of genes related to B cell survival and proliferation (IL10, IFR4, MYD88, JAK1, STAT3, BCL2, CD72) ^40–42^, immune attraction (CCL22, CCR4)^43^, glycolysis (ENO2, STMN1, PKM, CDK5R1)^29^ and motility (ACTG1, ACTG2, CD37)^44,45^ which are characteristics commonly associated with B cell malignancies (Figure 4A and 4C; Table S7). Notably, the cell-cell communication analysis based on ligand-receptor interactions reveals higher communication scores related to B cell immunosuppressive activities in Cell-Eco 1 (CD52, GZMB, IL16)^46–48^ (Figure 4B).

**Figure 4:**
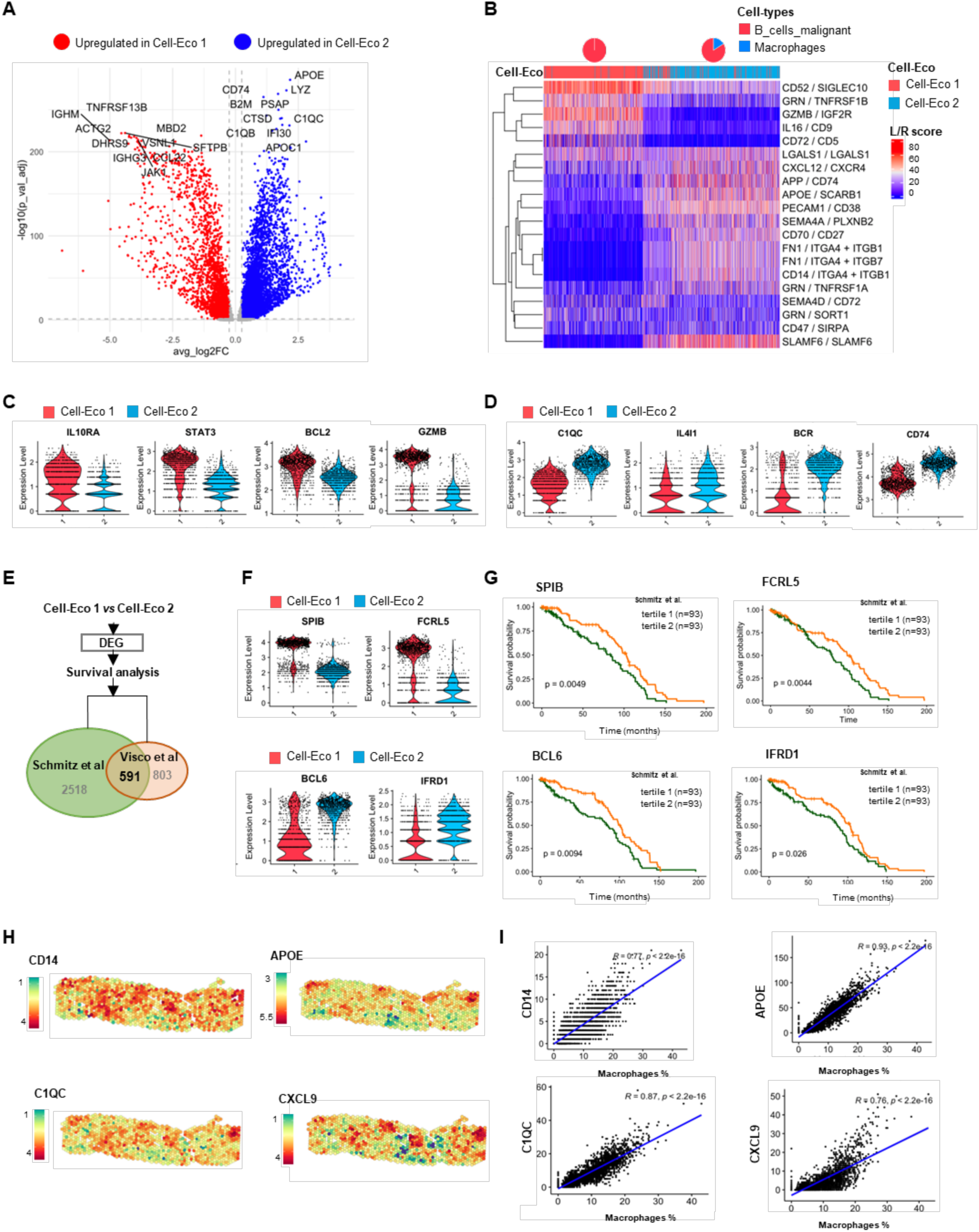
Tumor-supportive macrophages are the predominant cell type in direct contact with malignant B cells and shape the tumor microenvironment architecture. A) Volcano plot showing the DEGs between Cell-Eco 1 (in red) and Cell-Eco 2 (in blue), based on log2FC>0.25 and adjusted p-value<0.05. P-values were determined by the Wilcoxon-rank sum tests. Top 10 genes with lower adjusted p-values are annotated in the plot. B) Heatmap of ligand/receptor interaction scores based on intra-spot gene expression, between Cell-Eco1 (mean malignant B cell=100%) and Cell-Eco 2 (mean malignant B cells= 83.68, mean macrophages= 16.32%). Equations describing the score calculations are provided in the methods. C-D) Violin plots showing genes differentially upregulated in Cell-Eco 1 (C) and Cell-Eco 2 (D) (log2FC>0.25 and adjusted p-value<0.05 from the Wilcoxon-rank sum tests). E) Venn diagram illustrating the intersection of significant genes for survival analysis in two datasets of RNAseq DLBCL. Genes were considered significant if the Log-rank test p-value was below 0.05. The input for the survival analysis was DEGs between Cell-Eco 1 and Cell-Eco 2 (log2FC>0.25 and adjusted p-value<0.05 from the Wilcoxon-rank sum tests). F) Violin plots showing genes differentially upregulated in Cell-Eco and Cell-Eco 2 and significant for survival analysis in two independent RNAseq DLBCL datasets. G) Kaplan-Meier curves with Long-rank test p-value showing overall survival of patients divided by tertiles 1 and 2 for specific genes expression. H) Spatial representation of gene expression organization of spatially auto-correlated genes on a representative ST DLBCL sample. I) Scatter plots representing correlations between gene expression of spatially autocorrelated genes and the percentage of macrophages deconvoluted per spot in low infiltrated group of DLBCL ST samples. P values from pearson correlation.

Consistent with the presence of Macrophages in Cell-Eco 2, this Cell-Eco exhibits an upregulation macrophage lineage genes (CD163, CD68 and CD14), together with markers of activation, inflammation and antigen presentation (CD38, LYZ, STAT1, PSAP, CSTB, CD74, BCR)^49,50,51,52,53^. Interestingly, Cell-Eco 2 spots were highly enriched in genes associated to the C1q+ macrophage phenotype (C1QC, C1QA, C1QB, APOE, FOLR2, TREM2 and MRC1)^54^, when compared to Cell-Eco1, suggesting active cell-cell communication between malignant B cells and macrophages (Figure 4A, 4B and 4D; Table S7).

Pathway enrichment analysis based on differentially expressed genes between Cell-Eco 1 and Cell-Eco 2 revealed that Cell-Eco 2 are characterized by a macrophage and activation of the complement signatures (Figure S5B). Similarly, IL4I1 marker, associated to anti-inflammatory M2 polarization, was identified as upregulated in Cell-Eco2 (Figure 4D; Table S7). This phenotype is supported by the intra-spot analysis of communication molecules based on ligand-receptor expression levels. We identified distinct communication axes involved in the downregulation of macrophage anti-tumor functions, such as LGASLS1-LGASLS1^55^,, antigen presentation like APP-CD74, and immune cellular lymph node homing axis (CXCL12-CCR4)^56^, a pathway that has been implicated in TAMs infiltration (Figure 4B). The infiltration of the macrophages into the malignant microenvironment was confirmed by applying module scores of the germinal center (GC) vs inter-follicular (IF) macrophages signatures (MacroSig) from Lui et al^19^. The macrophages from Cell-Eco2 present a higher score of GC signatures compared to lymph node control macrophages, which exhibit a higher IF score as expected (Figure S5C).

To investigate the prognostic potential of differentially expressed genes between Cell-Eco 1 and Cell-Eco 2, we conducted a survival analysis on external RNA-seq datasets. We identified 591 genes that shows survival significance in two independent datasets: Schmitz et al.^27^ and Visco et al.^28^ (Table S8, Figure 4E). We found genes upregulated in Cell-Eco 1 and associated with worse prognosis like SPIB, a transcription factor of the PI3K-AKT pathway with anti-apoptotic effect in DLBCL^57^; FCRL5, a surface marker upregulated in malignant cells for which clinical therapies are investigated^58^; MALT1, a component of the NF-κB chronic signaling in DLBCL^59^; and JAK3, a member of the Janus kinase required for cell survival in DLBCL^60^ (Figure 4F and 4G; Figure S5D and S5E). On the other hand, among the genes with higher expression in Cell-Eco 2 and related to worse prognosis, we noted the protooncogene BLC6, a predictor of DLBCL poor survival^61^; IFRD1 a metabolic regulator implicated in autophagy inhibition^62^, and TP53I13 which promotes metastasis in glioma^60,63^ (Figure 4F and 4G; Figure S5D and S5E).

Subsequently, we performed spatial autocorrelation analysis, to identify the genes with similar expression levels in spatially proximate spots within the tissue^64^. A total of 210 genes were spatially autocorrelated in the low infiltration group of samples (Table S9). Interestingly, 39 of these genes show positive and significant correlation (R>0.5, p<0.05) with the frequency of deconvoluted macrophages while none of the spatially restricted genes were correlated with malignant B cells in this group (Figure 4H and 4I; Figure S5F, Table S9). This suggests that tumor-associated macrophages may play a role in the spatial organization and cellular remodeling of the TME in DLBCL, potentially contributing to the exclusion of other immune cells and enabling the immune evasion of malignant B cells.

## Discussion

Next generation sequencing (NGS) analysis of the tumor microenvironment (TME) in DLBCL has shown a link between immune cell infiltration and patient prognosis^10,15,27,65^. However, integrating spatial information with large-scale profiling of the immune TME in DLBCL has been limited due to the diffuse nature of the tissue^17,19, 29^. This study overcomes these challenges by using spatial transcriptomics (ST) to uncover spatially-restricted gene expression profiles within DLBCL tissues.

While, spatially defined composite cell neighborhoods (CNT) have been previously described by Wright et al^17^ using a 27-plex Multiplex Ion Beam Imaging (MIBI) panel, our study employed unsupervised clustering, independently of predefined panels. Our analysis identified six distinct immune cellular ecosystems (Cell-Eco) within the DLBCL TME, each defined by unique cellular composition, communication patterns and neighborhood characteristics. These Cell-Eco not only reflect the diverse immune cell types present in DLBCL tissues but also allow for the classification based on immune cell infiltrate. Indeed, in line with Wright et al findings, we observed that Cell-Eco varied in proportions of tumor and immune cells, that were inversely correlated. Importantly, we demonstrated that these Cell-Eco have significant prognostic potential, as their gene signatures stratify DLBCL patients by overall survival (OS) in independent datasets^27,28^, emphasizing the need for Cell-Eco-specific analyses to understand their biological and clinical significance.

One notable finding is the characterization of Cell Eco 4, which is abundant in highly infiltrated samples and characterized by a significant proportion of CD4+ T cells. The clinical relevance of CD4+ T cells has been well-documented, particularly in patients treated with R-CHOP, where their presence along with myofibroblasts (MFs) and dendritic cells (DCs), was associated with improved survival post-treatment^12,13^. In Cell-Eco 4, CD4+ T cells share coordinates with non-malignant B cells and a low proportion of malignant B cells. This pattern could reflect a reminiscence of healthy lymph node tissue. Interestingly, higher Cell-Eco 4 signature score associated with better OS in the Schmitz dataset^27^. This suggests that CD4+ T cells, even within varying colocalization profiles, play a crucial role in influencing patient clinical outcome.

Conventional dendritic cells (cDCs), which are key players in antigen presentation, were detected only in Cell-Eco 6, the ecosystem with the most diverse immune cell infiltration. Contrary to previous study from Wright et al.^17^, suggesting that DCs and macrophages are mutually exclusive, our findings reveal their intra-spot co-occurrence, challenging existing paradigms. Notably, cDCs in Cell-Eco 6 do not directly colocalize with malignant B cells but rather with other immune cells, suggesting that cDCs may contribute to immune cell recruitment or immune response coordination.

A remarkable result of our ST analysis derives from the comparison between Cell-Eco 3 and Cell-Eco 5, both containing CD8+ T cells and macrophages but differing in the malignant status of B cells. Despite similar cell composition, these ecosystems exhibit distinct transcriptomic profiles with opposite impacts on patient OS. For instance, Cell-Eco 3 (containing malignant B cells) is associated with poorer prognosis and exhibits higher expression of CCL22, a chemokine involved in Tregs recrutement^66^ and immune evasion in various cancer types, including DLBCL^67,68^. In addition, CCL22 expressed by M2 macrophages has been associated with metastasis^69^ and drug resistance^70^. These findings support an important role of CCL22 in the TME, which emerges as a potential clinical target for DLBCL patients with high Cell-Eco 3 signatures.

Another interesting finding is the upregulation CD24/SIGLEC10 communication axis in Cell-Eco 3 compared to Cell-Eco 5. While monoclonal antibody therapy targeting CD24 in DLBCL have shown limited efficacy^71,72^, our results suggest that targeting this axis should be restricted to the patients presenting high score of Cell-Eco 3.

In addition, we observed that Cell-Eco 3 presents a high glycolytic profile coupled with upregulation of anti-inflammatory signaling like STAT3. STAT3 is a well-studied component of the JAK/STAT signaling pathway involved in lymphomagenesis and tumor progression^60^. This pathway was shown to be constitutively activated in B-cell-like (ABC) DLBCL cells^41^. Interestingly, a recent study by “Dai et al.”^29^ identified clusters of malignant B cells and TAM (Tumor Associated Macrophages) presenting a similar glycolytic profile within an exhausted immune microenvironment. This cluster of cells was associated with poor prognosis in DLBCL.

Conversely, Cell-Eco 5 (composed by CD8+ T cells, macrophages and non-malignant B cells) was associated with better OS in two independent RNA-seq datasets^27,28^. Compared to Cell-Eco 3, Cell-Eco 5 displays higher expression of genes involved in lymph-node immune cell trafficking (CXCL12, CCL21 and CCL19)^34^, T cell activation (CD40LG, CD200, LAX1, LAG3) ^35,37,73–75^ and memory differentiation (TCF7, IL7R)^76^. These findings underscore the clinical significance of assessing cellular ecosystem functional properties, beyond cell composition.

Our analysis also sheds light on the role of macrophages in DLBCL. In DLBCL, macrophages have been associated with poor clinical outcomes in several studies^77–79^. Recently, Liu et al.^19^ identified distinct macrophages signatures with prognostic significance in DLBCL. To characterize the macrophages in direct proximity to malignant B cells, we performed differential expression analysis (DEA) between Cell-Eco 1 and Cell-Eco 2, the two ecosystems with highest proportion of malignant B cells but differing only by the presence of macrophages in Cell-Eco 2.

This analysis revealed that Cell-Eco 1 (composed mainly of malignant B cells) expresses markers typically associated with B cell activation, proliferation and metastasis. Nevertheless, the higher expression of markers such as IL10, CD52, IL16 and GZMB reflects the immunosuppressive functions of malignant B cells in this Cell-Eco. Interestingly, GZMB-producing B cells have been described to suppress T cell proliferation and cytokine production in different cancer types and lymphoma cell lines^80,81^. Thus, GZMB-producing malignant B cells may contribute to T cell exclusion in ecosystems rich in malignant B cells.

In contrast, Cell-Eco 2 (composed of malignant B cells together with macrophages) exhibits upregulation of macrophage lineage genes, particularly those associated with the C1q+ subpopulation^54^. This phenotype goes beyond M1/M2 dichotomy, suggesting a possible role of the complement pathway in macrophage polarization and tumor proliferation in this Cell-Eco. Interestingly, enrichment in complement recognition patterns by macrophages was associated with poorer survival in a recent study from Liu et al ^19^. Another study revealed that C1QB-expressing macrophages are the source of cholesterol efflux linked to CAR-T resistance in refractory DLBCL^82^. These findings underscore the significance of complement activation and metabolic pathways for macrophage activity, highlighting their potential role in DLBCL treatment response. Notably, Cell-Eco 2 was the most abundant in sample DLBCL_3, the only sample analyzed post-relapse from R-CHOP treatment. This suggests a shift in cellular composition and colocalization after treatment, with macrophage phenotype potentially serving as an indicator of treatment response.

Moreover, in samples with the highest proportions of malignant B cells, macrophages levels were correlated with the expression of spatially restricted genes. This implies that, despite the wide dispersion of malignant B cells, macrophages might be the key immune cells shaping the spatial organization within highly malignant microenvironments. Other genes with spatial restrictions were associated with extracellular matrix (ECM) rather than immune cells proportions. However, our study did not investigate the role of stromal cells, particularly cancer associated fibroblasts (CAFs) which, together with macrophages, could orchestrate immune cell infiltration and tissue architecture in the TME.

In conclusion, our spatial transcriptomic study expands the understanding of DLBCL tissue architecture, revealing novel cellular organizations, cell states and communication patterns organized into distinct cellular ecosystems. Importantly, the clinical relevance of these cellular ecosystems was validated by the stratification of patients based on overall survival using external public datasets. Beyond identifying cell type phenotypes, our study stresses the need for a deep characterization of cell-cell interactions and functional states to better understand the TME dynamics and clinical outcomes in DLBCL. Finally, this study provides a valuable data resource for further in-depth analyses of the TME in DLBCL and the identification of potential therapeutic targets or biomarkers.

### Limitations of the study

This study has several limitations including the small sample size and the limited coverage of the TME due to the small size of the tissue sections analyzed. While ST analysis across different biopsies from the same patients resulted in similar Cell-Eco patterns in our study, a larger sample size and tissue sections could enhance the robustness of our findings.

Another limitation is the sequencing resolution of Visium ST technology, which captures transcripts at a spot level, potentially missing rare cell subsets. The use of novel ST technologies with single-cell resolution or the development of spot segmentation methods could provide more precise insights into the spatial distribution and cellular interactions within the TME. Lastly, the absence of stromal cells transcriptomics signatures in the scRNA-seq dataset used for deconvolution may hinder complete tissue characterization^83^.

## METHODS

### Key resources table

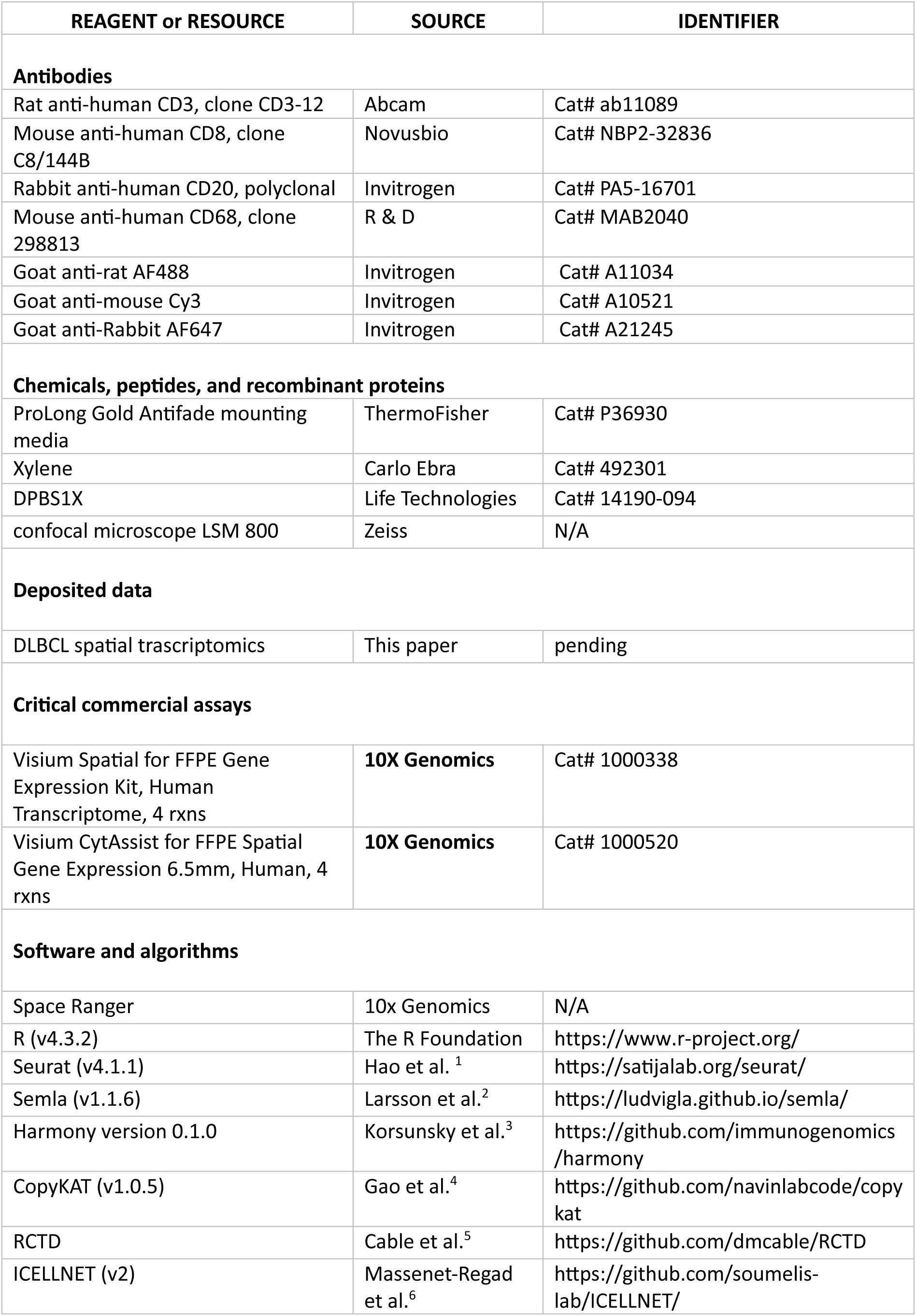
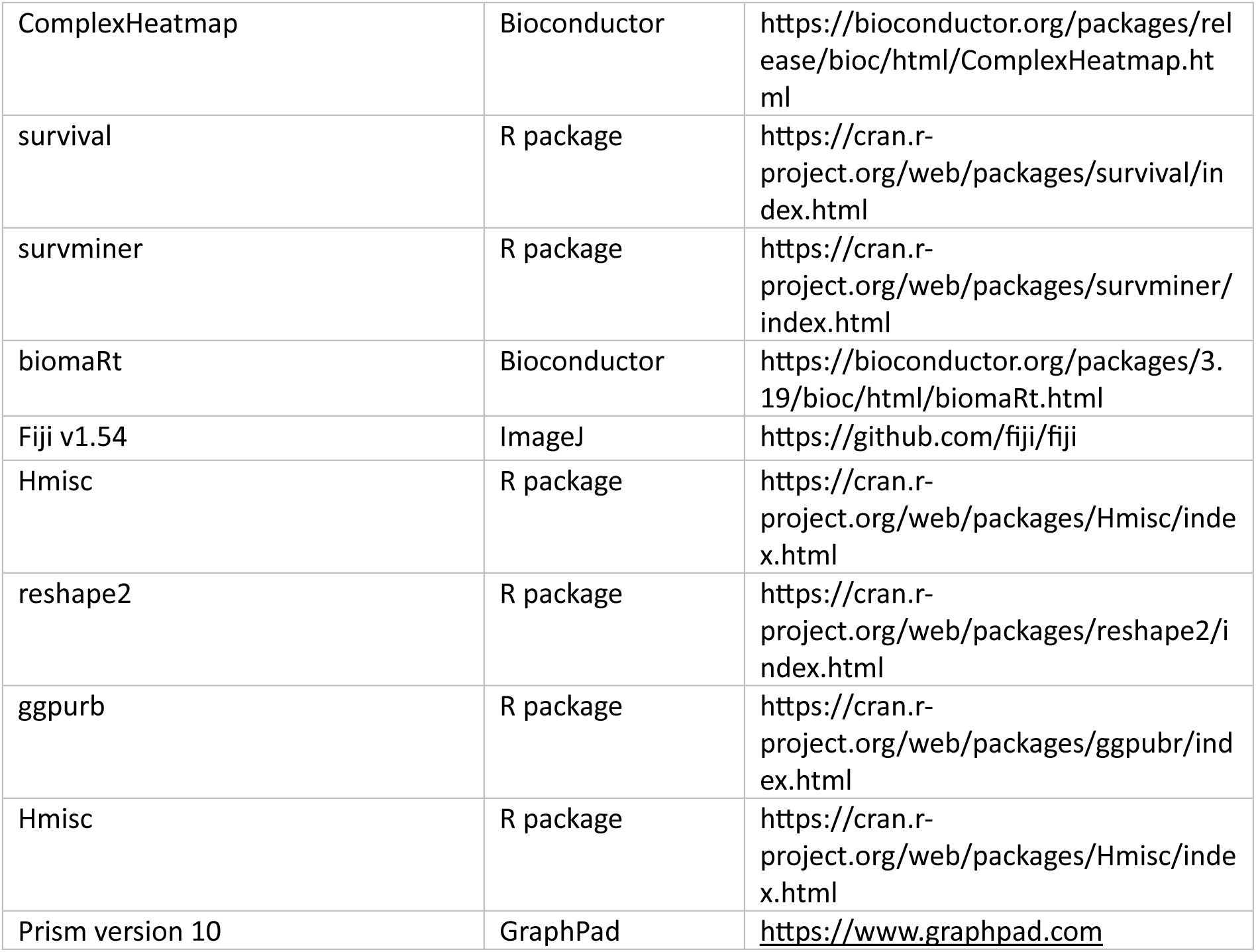

#### Lead contact

Further information and requests for resources and reagents should be directed to and will be fulfilled by the lead contact, Pierre Tonnerre.

#### Materials availability

This study did not generate new unique reagents.

#### Data and code availability

- Data: The spatial transcriptomic data generated for this study will be deposited in Gene Expression Omnibus (GEO) upon publication. The scRNA-seq data used for spatial data deconvolution was retrieved from heiDATA under accession code <u>VRJUNV</u> and from GEO under the accession number <u>GSE182436</u>. The public DLBCL expression array dataset was retrieved from GEO under the accession number <u>GSE31312</u>. DLBCL RNA-seq dataset was obtained through a data access request to the National Institutes of Health (NIH) database of Genotypes and Phenotypes (dbGaP) under the accession number: <u>phs001444.v2.p1</u>.
- Any additional data is available from the lead contact upon request.

## EXPERIMENTAL MODEL AND SUBJECT DETAILS

### Ethics statement and Human samples

All samples were obtained following the principles of Good Clinical Practice and the Declaration of Helsinki, ensuring compliance with ethical regulations.

Samples from DLBCL patients (n=8) were obtained from the Pathology Department at the Hospital Saint Louis (Paris, France). Patients were previously diagnosed by certified physicians in the Haemato-Oncology department of the Hospital Saint Louis (Paris, France) and the histopathological diagnosis was confirmed by an expert hematopathologist according to the international classifications. The clinical characteristics of patient samples are provided in Table S1. Non-metastatic lymph node samples (LN) and tonsil were obtained from individuals undergoing surgical procedures with no cancer indications at the Hospital Bichat and Robert Debré (Paris, France). Informed consent for tissue collection and research participation was obtained from each patient, without any financial compensation.

### Sample selection

All tissues were Formalin-fixed paraffin-embedded (FFPE) upon resection. DLBCL and Secondary Lymphoid Organ (SLO) samples were pre-selected based on the confirmed diagnosis and cell viability upon histopathology examination. Sample selection was dependent on the RNA quality control index determined by the proportion of RNA fragments over 200 nucleotides in length (DV200). Specifically, samples with a DV200 ≥ 30% were included in the study, in accordance with the recommendations and guidance from the 10X Genomics technical support team. The entire slides

## METHOD DETAILS

### Spatial transcriptome sequencing

Spatial transcriptomic (ST) data of DLBCL and SLO (reactive lymph node and tonsil) samples were generated in two independent experiments: 2 DLBCL samples and 2 SLO control data were generated from the Visium Spatial Gene Expression for FFPE with “Direct placement” and 8 DLBCL samples and 2 SLO controls data was generated using the CytAssist instrument or “CytAssist-enabled” assay. From each block, sections of 4 μm of thickness were printed to the Visium slides’ capture areas of 6.5 × 6.5 mm^2^ and stained with hematoxylin and eosin following the corresponding 10x Genomics Visium FFPE tissue processing protocol (CG000408 or CG000518). The entire section of DLBCL samples were placed on the Visium slides. Samples underwent deparaffination, imaging and decrosslinking (CG000409 or CG000520), followed by probe hybridation, and ligation, tissue removal, and library preparation following corresponding User Guides (CG000407 or CG000495). Each FFPE library was sequenced using a NovaSeq6000 Sequencing System (Illumina, CA, USA). FASTQ files were processed using the GRCh38 transcriptome data from 10× Genomics and the Space Ranger pipeline (v2.0.1).

### Spatial transcriptomics data processing

Spatial transcriptomic data of each sample were processed individually, employing R (v4.3.2) and Seurat (v4.1.1)^84^. Capture areas shared by multiple biopsies were segmented based on their coordinates with the package Semla (v1.1.6)^64^ to ensure one Seurat object per sample. Each object was processed individually; data were normalized and scaled with the *SCTransform* function to remove variation due to sequencing depth across spots. The high variable features were detected and the transformed data were stored and processed on SCT assay. To reduce dimensionality, principal component analysis (PCA) was applied to the 2000 most variable genes, employing the variance-stabilizing transformation approach.

### Control Tissue region annotation

For SLO control samples, unsupervised gene expression analysis was conducted at a spot level. Spots clustering was performed by computing the Louvain clustering algorithm^85^, applied to a shared nearest-neighbor (SNN) graph, using the functions *FindNeighbors* and FindClusters. *FindAllMarkers* was used to perform a Wilcoxon-rank sum test to identify differentially expressed genes between clusters. Genes were considered differently expressed if their |log2FC| was higher than 0.25 and their adjusted p-value below 0.05. Manual cluster annotation was performed based on the differentially expressed genes.

### Preprocessing and annotation of single-cell RNA sequencing public data

The scRNA-seq data analysis was conducted on R (v4.3.2) with the package Seurat (v4.1.1)^84^. The scRNA-seq of DLBCL, reactive lymph node (rLN) and tonsil samples were collected from Roider et al.^14^, Steen et al.^15^ and an internal scRNA-seq dataset. Supplementary information about scRNA-seq samples is provided in Table S9. Each sample data was pre-processed individually to conserve cells expressing a minimum of 200 genes and less than 10% of mitochondrial genes. The data were then normalized based on library size, followed by a log2 transformation. Dimensionality reduction was performed by PCA as previously described. Genes identifiers were homogenized with the package bioMart^86^ across datasets and samples to the GRCh38 reference: GENCODE v32/Ensembl 98^87^ annotation.

Subsequently, the data was merged into two sets, DLBCL or SLO, normalized for library size and log2-transformed. Samples of each set were integrated based on their first 50 Principal Components (PC), using the package Harmony^88^ (v0.1.0). Cell clustering was performed as previously described for ST data. Malignant B cells were annotated based on their chromosomal copy number variation with the R package CopyKAT (v1.0.5)^89^. The identity of each cluster was determined through manual inspection and prior knowledge of cluster-specific markers (p-values<0.05 and |log2FC|>0.25) identified within the output list of a Wilcoxon-rank sum test. Top differentially expressed genes for cluster annotations are provided in Table S2.

### Single-cell RNA sequencing and Spatial Transcriptomic data integration

Following the pre-processing of ST data, genes were filtered to retain common genes between assays. The spots of each sample were decomposed using the corresponding public scRNA-seq dataset (either DLBCL or SLO) as reference. The cell type proportions per spot were generated using the probabilistic method RCTD^20^ (Robust Cell Type Decomposition) on *multi-mode*. The RCTD algorithm was applied to the gene sets previously homogenized across samples that were analyzed using both V1 and V2 Visium assays (Figure S5G). All genes selected from the scRNA-seq data for cell type decomposition were included in both assays for all samples (Figure S5H).

### Spot clustering based on cell type proportion

First, a matrix of cell-type proportions was constructed by merging the individual RCTD outputs of each DLBCL ST sample. Cell type proportions lower than 10% were filtered and considered as not biologically meaningful. After filtration, spots were scaled to ensure a cumulative sum of 100% of cell proportions. Leiden algorithm^26^, a graph-based clustering method implemented by Python v3.x, was applied in R to identify groups of spots based on similar cell composition. Cluster granularity was set to a resolution of 0.2 since a different resolution did not improve the biological interpretation of the clusters. Spots were labeled based on the Leiden clustering results as Cellular ecosystems (Cell-Eco). Cell-type means higher than 10% were considered biologically significant for the Cell-Eco identities. Spots were filtered and rescaled based on cell-types means per Cell-Eco as previously described.

### Differential expression analysis and module scores of Cell-Eco

DLBCL ST samples from the V2 assay were merged into one Seurat object, then normalized and scaled for counts variance stabilization with the *SCTransform* function of Seurat. Differential expression analysis (DEA) based on the non-parametric Wilcoxon rank sum test was performed between the Cell-Eco labels, transferred from the RTCD output to their corresponding spot in the SCT data matrix. DEA was carried out across all the Cell-Eco by the Seurat function *FindAllMarkers* while *FindMarkers* conducted the DEA between two specific Cell-Eco. Pathway enrichment analysis was performed on DEG using Metascape^90^ and/or DisGeNET^91^ when necessary.

Module scores (https://satijalab.org/seurat/reference/addmodulescore) from publicly available gene signatures of interest ^19^ were applied to the ST data in order to compare relative expression of gene signatures between Cell-Eco. For the comparison of module scores between Cell-Eco 2 and control samples, spots from the rLN sample of the V2 assay were filtered to retain those with macrophages proportions higher than 10%.

### Neighbor scores

Considering a maximum of six neighbors per spot, we applied the function *RegionNeighbors* of the package Semla to extract Cellular ecosystem’s identity of spots spatially co-localizing. The neighbor score (N score) was calculated as a frequency of neighbor interactions by the following formula:

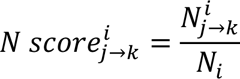

Where *j* is a Cell-Eco identity spatially located next to another unique identity, *k,* in sample *i*. Therefore, *N_j→k_* refers to the number of Cell-Eco *j* neighbor to a Cell-Eco *k* in a sample *i*, divided by the total neighbors of the sample (*N_i_*). The neighbor score of each group of samples was calculated as a mean of neighbor scores as follows:

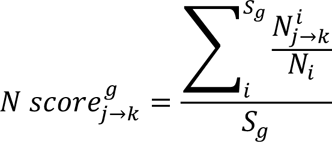

Where *g* is the identity of the group of samples, and *S* the number of samples. Consequently, the neighbor score of a Cell Eco *j* next to a Cell eco *k* per group *g* is the sum of neighbor scores of each sample *i* belonging to the group *g* divided by total number of samples in the group *S_g_*.

### Intra-spot ligand-receptor communication

Ligand-receptor interactions were assessed by scoring their expression level using the frameworks based on databases of known receptor-ligand interactions ICELLNET(v2)^92^. The communication scores (C score) were calculated per spot as follows:

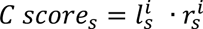

where 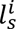 corresponds to the average expression of a ligand *i* in the spot *s* and 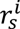 refers to the average expression of the corresponding receptors to ligand *i* in the same spot *s*.

### Survival analysis

The signature for each Cell-Eco was defined as the percentile 5 of differentially expressed genes (p<0.05 and |log2FC|>0.25) ordered by adjusted p-value. These signatures were applied to two different DLBCL bulk gene expression cohorts,^27,28^ to investigate their potential implication in patients’ survival. For each Cell-Eco signature, a score previously used in the literature by Liu et al.^19^ was calculated as follows:

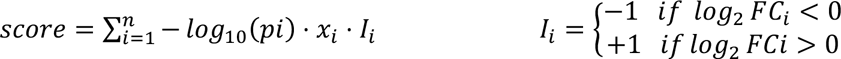

where *n* is the number of genes in a signature, *pi* is the adjusted p-value of the gene *i*, previously calculated by Wilcoxon rank sum test, *xi* is the expression of the gene *i* in the bulk matrix. *I* corresponds to the sign of the log_2_FC value; positive if the gene *i* is upregulated and negative if its downregulated for a specific Cell-Eco signature. Signature scores for each Cellular ecosystem were scaled and centered prior to the hierarchical clustering for patient stratification.

For the individual Cell-Eco signatures survival assessment; patients were divided into three groups based on their signature scores, using tertiles^93^ as the cutoffs. Tertile 1 included patients with higher Cell-Eco signature scores, tercile 2 included patients with intermediate signature scores, and tertile 2 included patients with lower signature scores. Alongside, survival analysis was applied as previously described, to the entire list of differentially expressed genes (p<0.05 and |log2FC|>0.25) between pairs of Cell-Eco, in order to identify particular genes with potential prognostic power.

Survival analysis was conducted with the R packages survival^94^ (v3.6-4) and survimer (v0.4.9). Kaplan-Meier^95^ curves of overall survival (OS) among stratified patient groups and for one Cell-Eco signature between tercile groups of patients were compared using the long-rank test.

### Hierarchical clustering

Hierarchical clustering of communication molecules and patient stratification was performed based on Euclidean distances with the R package ComplexHeatmap (v2.20.0). The heatmaps dendrogram were built under the ward.D2 aggregation method^96^.For patient stratification scores were centered and scaled prior to clustering.

### Spatial autocorrelation

Spatially autocorrelated genes were identified using the CorSpatialFeatures function of Semla package^64^. Only spatial correlation scores greater than 0.5 were considered for the gene selection, since these genes show tendency of similar expression in proximate spots of the tissue.

### Immunofluorescence staining and image analysis

The samples were first deparaffinized by heating the slides to 56°C and the tissue was rehydrated by sequentially immersing the slides in the following solutions for 5 minutes: xylene (2 washes), 100% ethanol, 75% ethanol, 50% ethanol and deionized water (this last one for 10 minutes). Then, the samples were immersed in the prewarmed antigen retrieval solution pH 9 at 95°C in a water bath for 20 minutes and allowed to cool at room temperature (RT). The samples were washed with deionized water for 10 minutes at RT and permeabilized by incubating the slide in deionized water with 0.5% Triton. The slides were washed with water and PBS for 10 minutes. The edges of the slide were wiped, and the tissue was circled with the hydrophobic ink of the Dakopen. Next, samples were incubated with the relevant primary antibodies diluted in BFG buffer (BSA 2.5%, fish gelatin 0,1%, Glycine 100 mM, pH7) overnight at 4°C. Samples were washed three times with PBS for 5 minutes each wash and the relevant secondary antibodies were added diluted in BFG for 2 hours at RT. Slides were washed again three times in PBS for 5 minutes and were incubated with DAPI dye for 5 minutes. Finally, samples were washed 3 times with PBS, and slides were mounted by adding two drops of ProLong Gold Antifade mounting media on the coverslip. Images of the slides were taken in a confocal microscope LSM 800 (Zeiss). Images were analyzed on Fiji v1.54 (ImageJ2). The percentage of CD3+, CD8+, CD20+, or CD68+ cells in each sample was calculated by counting the cells stained and relative to the total number of cells corresponding to the DAPI staining. Briefly, the images were converted to binary images by adjusting the threshold approximately to 70% above and 0% below for DAPI staining and to 0% above and 10% below for antibody staining. Next, cells were segmented using the StarDist 2D plugin (doi:10.1007/978-3-030-00934-2_30) using the default values. Only particles with an area between 30 and 600 px2 were counted.

### Quantification and statistical analysis

Statistical analyses were conducted on R (v4.1.1) and GraphPad Prism (v10) software. Pearson correlation analysis was performed with the R package Hmisc. Differences in cell type proportions between groups were assessed by the non-parametric Kruskal-Wallis test for multiple comparison combined with a Dunn’s pot-hoc test for pairwise comparison. The R packages reshape2 and ggpubr were used for data analysis and visualization purposes.

The Wilcoxon-rank sum test with Bonferroni analysis was applied to identify differentially expressed genes between groups. Survival analysis was performed by utilizing Kapan‒Meier curves with a log rank test for significance. P-values or adjusted p-values below 0.05 (<0.05) were considered statistically significant for every test. The figure legends provide information about all statistical tests and parameters applied.

## Supplemental information

Supplemental information can be found online.

## Acknowledgments

This work was supported by the laboratory of excellence (Labex) Inflamex, the Fondation ARC pour la Recherche sur le Cancer (Sys-Can ARCPGA12021 020003101_358 7), and Institut National de la Santé et de la Recherche Médicale (INSERM) under the grant CHEMOTAXIS IN CANCER. We thank the Institut Curie PATHEX and NGS facilities, in particular Renaud Leclere, and Mylene Bohec for their technical support. We also thank the Institut de Recherche Saint-Louis imaging facility, in particular Niclas Setterblad, for their technical support. A.D.H was supported by a PhD fellowship from Servier. L.M.R. was supported by a PhD fellowship from La Ligue Contre le Cancer, and Fondation ARC. P.T. is supported by the ATIP-Avenir program from INSERM and CNRS.

## Authors contributions

A.D.H. conceived the project, build the circuit of samples, analyzed and interpreted the ST data, and wrote the manuscript. H.P.P. performed the tissue staining and quantification of immunofluorescence images and contributed to the manuscript writing. J.L.C. conducted quality control of the RNA from FFPE tissue sections. L.M.R. and M.V. gave technical and conceptual advice for bioinformatic analyses and contributed to the scRNA-seq analysis. J.C., C.C., V.M. and C.T. facilitated access to histological samples, clinical information and interpretation. V.B. and V.S. contributed to the study design, project management and funding. P.T. contributed to the study design, project management and funding, data interpretation, and supervised the work. All authors reviewed the manuscript.

