## Supplementary Figures for "Spatial transcriptomics unveils immune cellular ecosystems associated with patient survival in diffuse large B-cell lymphoma"

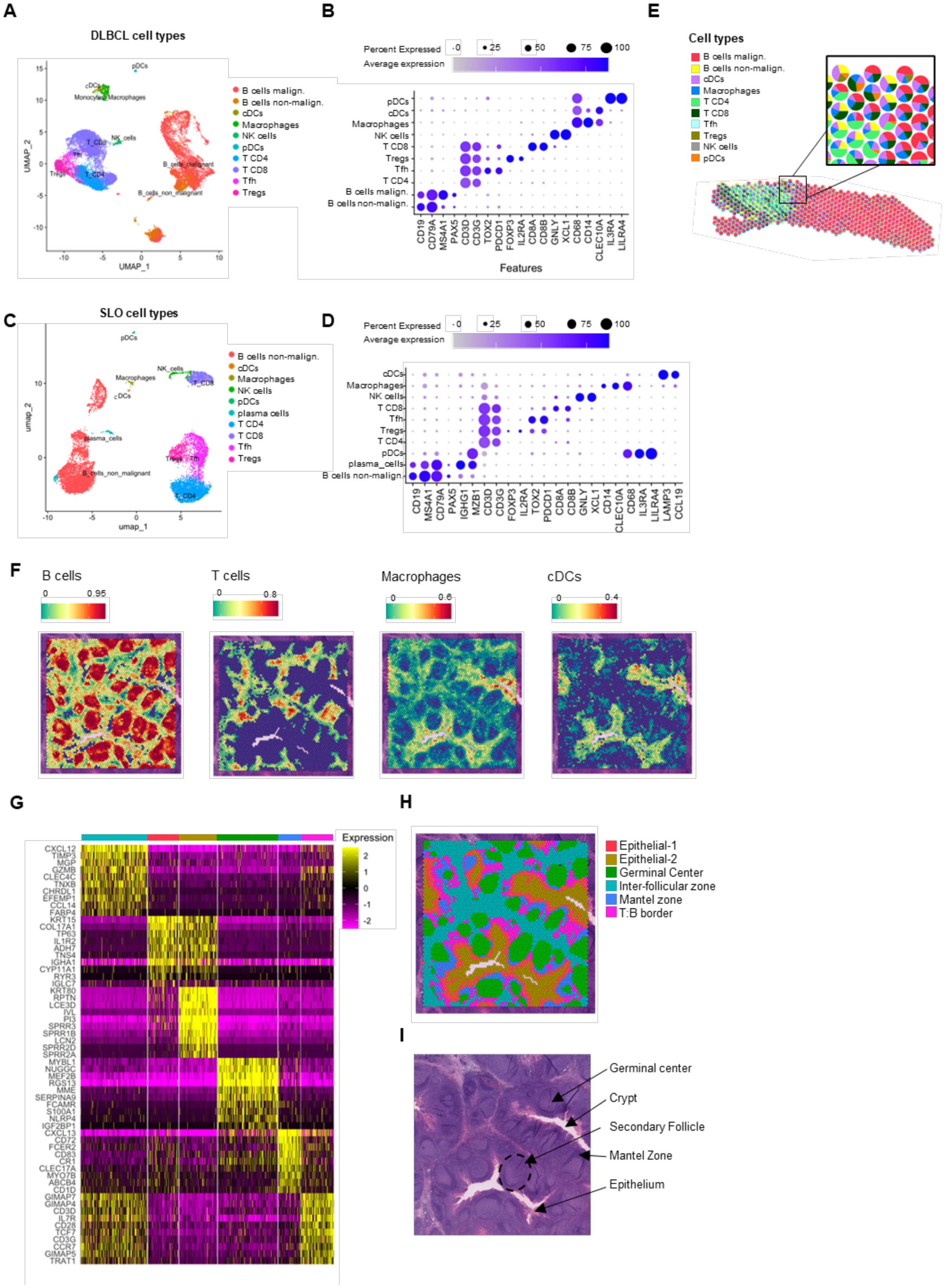

A
DLBCL cell types
pDCs
cDCs
Monocytes Macrophages
T_CD8
NK_cells
Tfh
Tregs
T_CD4
B_cells_malignant
B_cells_non_malignant
UMAP_1
UMAP_2
B cells malign.
B cells non-malign.
cDCs
Macrophages
NK cells
pDCs
T CD4
T CD8
Tfh
Tregs
B
Percent Expressed
0
25
50
75
100
Average expression
pDCs
cDCs
Macrophages
NK cells
T CD8
Tregs
Tfh
T CD4
B cells malign.
B cells non-malign.
CD19
CD79A
MS4A1
PAX5
CD3D
CD3G
TOX2
PDCD1
FOXP3
IL2RA
CD8A
CD8B
GNLY
XCL1
CD68
CD14
CLEC10A
IL3RA
LILRA4
Features
E
Cell types
B cells malign.
B cells non-malign.
cDCs
Macrophages
T CD4
T CD8
Tfh
Tregs
NK cells
pDCs
C
SLO cell types
pDCs
NK_cells
Macrophages
cDCs
T_CD8
plasma_cells
Tregs
Tfh
B_cells_non_malignant
T_CD4
umap_1
umap_2
B cells non-malign.
cDCs
Macrophages
NK cells
pDCs
plasma cells
T CD4
T CD8
Tfh
Tregs
D
Percent Expressed
0
25
50
75
100
Average expression
cDCs
Macrophages
NK cells
T CD8
Tfh
Tregs
T CD4
pDCs
plasma_cells
B cells non-malign.
CD19
MS4A1
CD79A
PAX5
IGHG1
MZB1
CD3D
CD3G
FOXP3
IL2RA
TOX2
PDCD1
CD8A
CD8B
GNLY
XCL1
CD14
CLEC10A
CD68
IL3RA
LILRA4
LAMP3
CCL19
F
B cells
0
0.95
T cells
0
0.8
Macrophages
0
0.6
cDCs
0
0.4
G
Expression
2
1
0
-1
-2
CXCL12
TIMP3
MGP
GZMB
CLEC4C
TNXB
CHRDL1
EFEMP1
CCL14
FABP4
KRT15
COL17A1
TP63
IL1R2
ADH7
TNS4
IGHA1
CYP11A1
RYR3
IGLC7
KRT80
RPTN
LCE3D
IVL
PI3
SPRR3
SPRR1B
LCN2
SPRR2D
SPRR2A
MYBL1
NUGGC
MEF2B
RGS13
MME
SERPINA9
FCAMR
S100A1
NLRP4
IGF2BP1
CXCL13
CD72
FCER2
CD83
CR1
CLEC17A
MYO7B
ABCB4
CD1D
GIMAP7
GIMAP4
CD3D
IL7R
CD28
TCF7
CD3G
CCR7
GIMAP5
TRAT1
H
Epithelial-1
Epithelial-2
Germinal Center
Inter-follicular zone
Mantel zone
T:B border
I
Germinal center
Crypt
Secondary Follicle
Mantel Zone
Epithelium

**Figure S1: RCTD deconvolution. Related to Figure 1.**

A) Single cell RNA sequencing public data, manually annotated. Uniform Manifold Approximation and Projection (UMAP) of 21990 single immune cells from 6 DLBCL samples. The UMAP show the annotation of 10 immune cell types, including malignant B cells, non-malignant B cells, dendritic cells (DCs), Macrophages, natural killer cells (NK cells), plasmacytoid dendritic cells (pDCs), CD4+ T cells (T CD4), CD8+ T cells (T CD8), follicular T helpers (Tfh), and regulatory T cells (Tregs). B) Gene expression Dot plot of main markers associated to each cell type annotated in Figure S1A. C) Single cell RNA sequencing public data, manually annotated. Uniform Manifold Approximation and Projection (UMAP) of 11108 CD45+ single cells from 4 SLO samples (1 tonsil and 3 reactive lymph nodes). The UMAP show the annotation of 10 cell types. D) Gene expression of main markers associated to each cell type annotated in Figure S1C. E) Spatial scatter pie plot of a representative ST DLBCL sample, showing deconvoluted cell type proportions per spot. F) Spatial representation of deconvoluted cell-types proportions on a tonsil section. G) Heatmap of the top 15 genes per cluster of a representative ST human tonsil sample. H) Spatial representation of annotated clusters of a representative ST human tonsil sample. I) Hematoxylin and eosin-stained slide of a representative human tonsil histologically annotated.

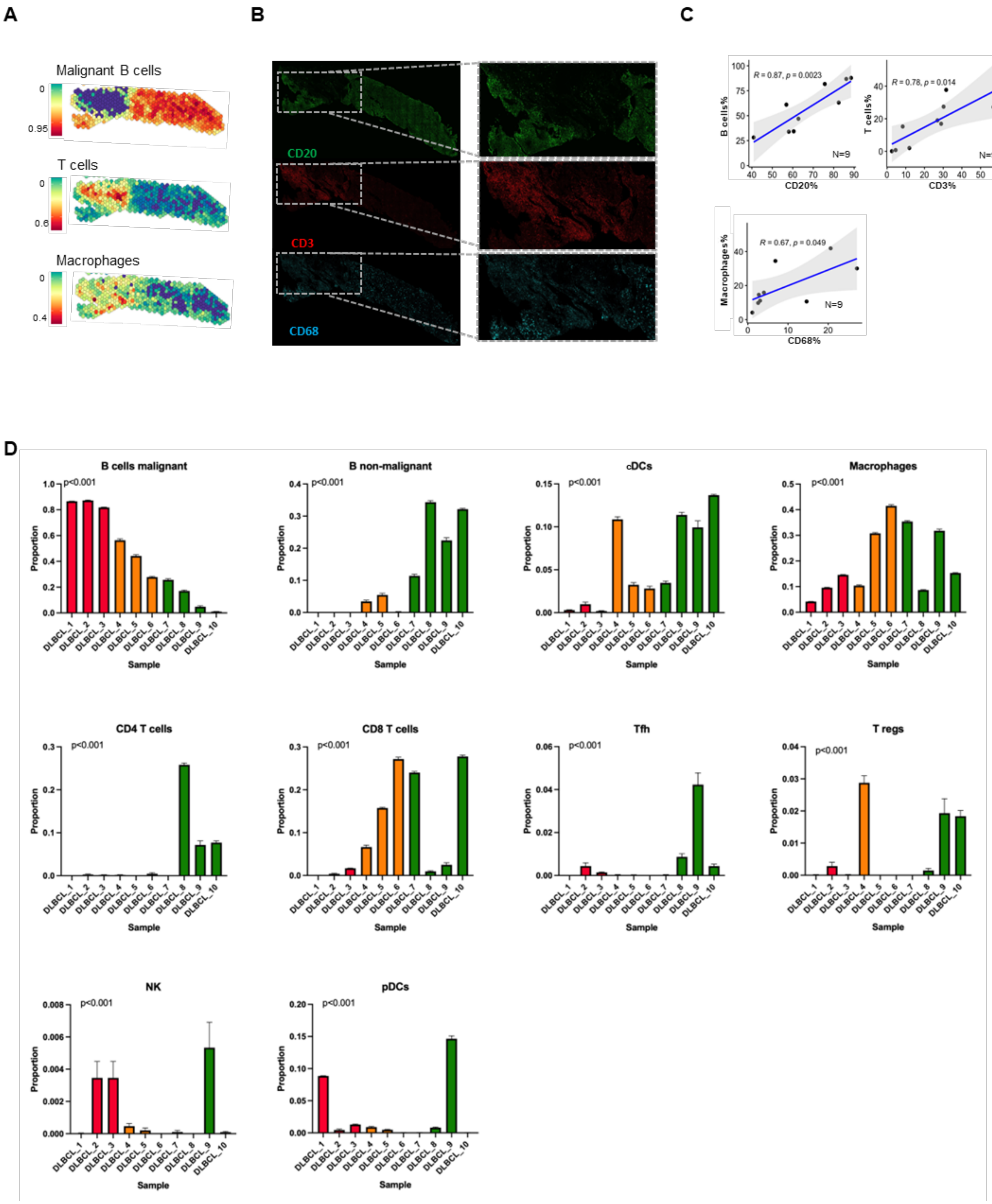
A
Malignant B cells
T cells
Macrophages
B
CD20
CD3
CD68
C
R = 0.87, p = 0.0023
N=9
B cells%
CD20%
R = 0.78, p = 0.014
T cells%
CD3%
R = 0.67, p = 0.049
Macrophages%
CD68%
D
B cells malignant
B non-malignant
cDCs
Macrophages
CD4 T cells
CD8 T cells
Tfh
T regs
NK
pDCs
p<0.001
Proportion
Sample
DLBCL_1
DLBCL_2
DLBCL_3
DLBCL_4
DLBCL_5
DLBCL_6
DLBCL_7
DLBCL_8
DLBCL_9
DLBCL_10

**Figure S2: Protein validation of RCTD deconvolution and RCTD-derived immune cell-type proportions across DLBCL ST samples. Related to Figure 1**

A) Spatial representation deconvoluted immune cell-types proportions (B cells malignant, T cells and Macrophages) in a representative ST DLBCL sample. B) Immune florescence images of selected markers CD20 (green), CD3 (red) and CD68 (cyan) from a representative ST DLBCL sample**.** C) Scatter plots illustrating the Pearson correlation between the proportions of deconvoluted cell types, and the proportion of cells positive for immunofluorescence staining for the corresponding marker. Each spot of the scatter plots represents one individual sample. D) Bar plot representing the mean proportions of RCTD-deconvoluted CD8+ T cells, CD4+ T cells, regulatory T cells, macrophages, natural killer cells, dendritic cells, plasmacytoid dendritic cells, dendritic cells, non-malignant B cells, malignant B cells. Values represent the mean proportion of cell-type ±SEM. Multiple comparisons were performed using Kruskal-Wallis statistical test and Dunn's post-hoc for pairwise comparisons (not shown).

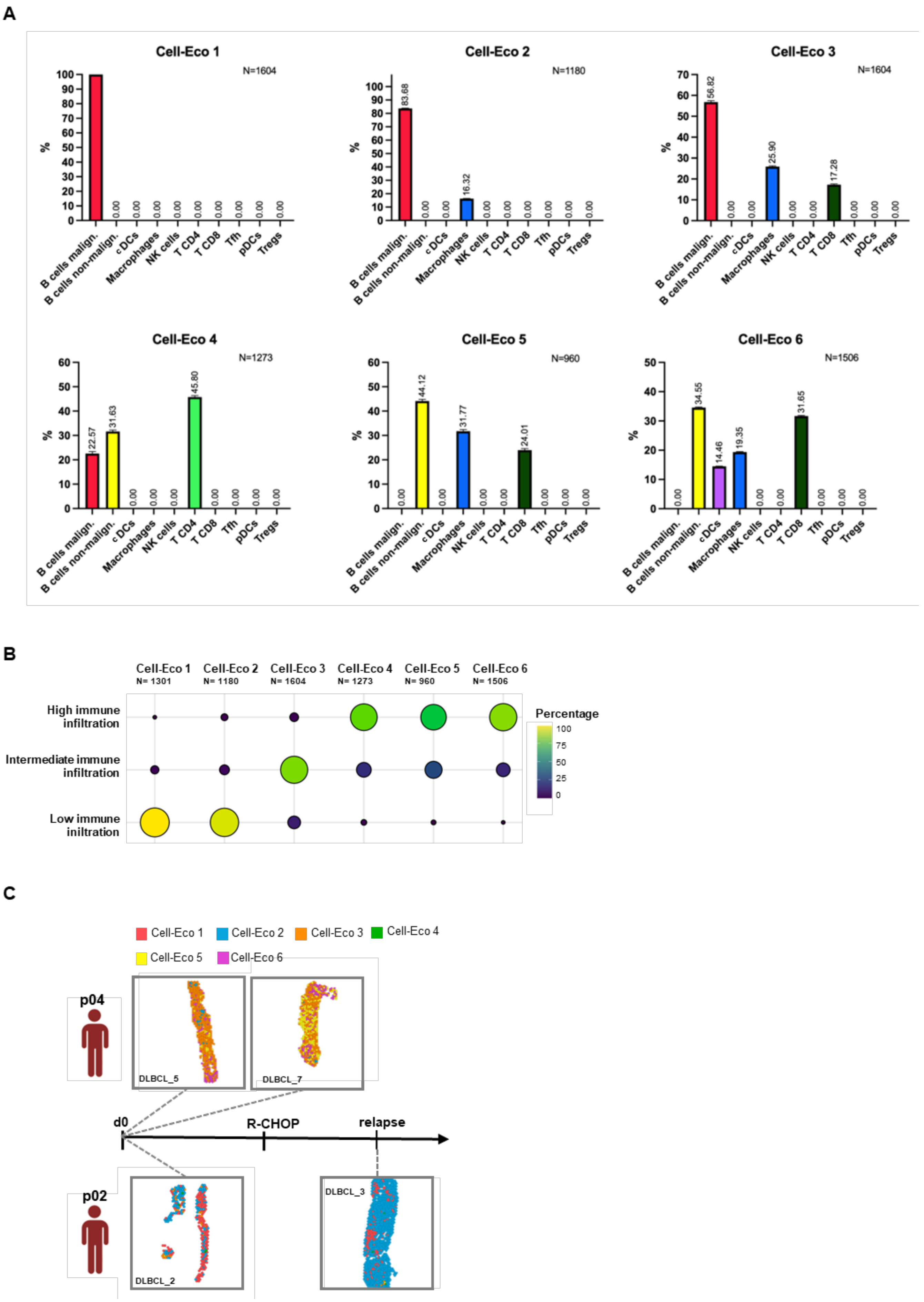
A
Cell-Eco 1
N=1604
%
100
0.00
Cell-Eco 2
N=1180
83.68
16.32
Cell-Eco 3
N=1604
56.82
25.90
17.28
Cell-Eco 4
N=1273
22.57
31.63
45.80
Cell-Eco 5
N=960
44.12
31.77
24.01
Cell-Eco 6
N=1506
34.55
14.46
19.35
31.65
B cells malign.
B cells non-malign.
cDCs
Macrophages
NK cells
T CD4
T CD8
Tfh
pDCs
Tregs
B
Cell-Eco 1 N= 1301
Cell-Eco 2 N= 1180
Cell-Eco 3 N= 1604
Cell-Eco 4 N= 1273
Cell-Eco 5 N= 960
Cell-Eco 6 N= 1506
High immune infiltration
Intermediate immune infiltration
Low immune iniltration
Percentage
100
75
50
25
0
C
Cell-Eco 1
Cell-Eco 2
Cell-Eco 3
Cell-Eco 4
Cell-Eco 5
Cell-Eco 6
p04
DLBCL_5
DLBCL_7
d0
R-CHOP
relapse
p02
DLBCL_2
DLBCL_3

**Figure S3: Cell-Eco-associated cell-type proportions and frequencies across sample groups. Related to Figure 2.**

A) Bar plots illustrating cell proportions filtered and scaled to 100% per Cell-Eco. Values represent the mean of cell-type proportion ±SEM. B) Frequencies of Cell-Ecos across sample groups as percentages. C) Spatial distribution of the Cellular Ecosystems in DLBCL ST samples biopsied from the same patient at different time points: baseline or pre-treatment (d0) and at relapse post-treatment (relapse). Distance between events do not reflect time lapse.

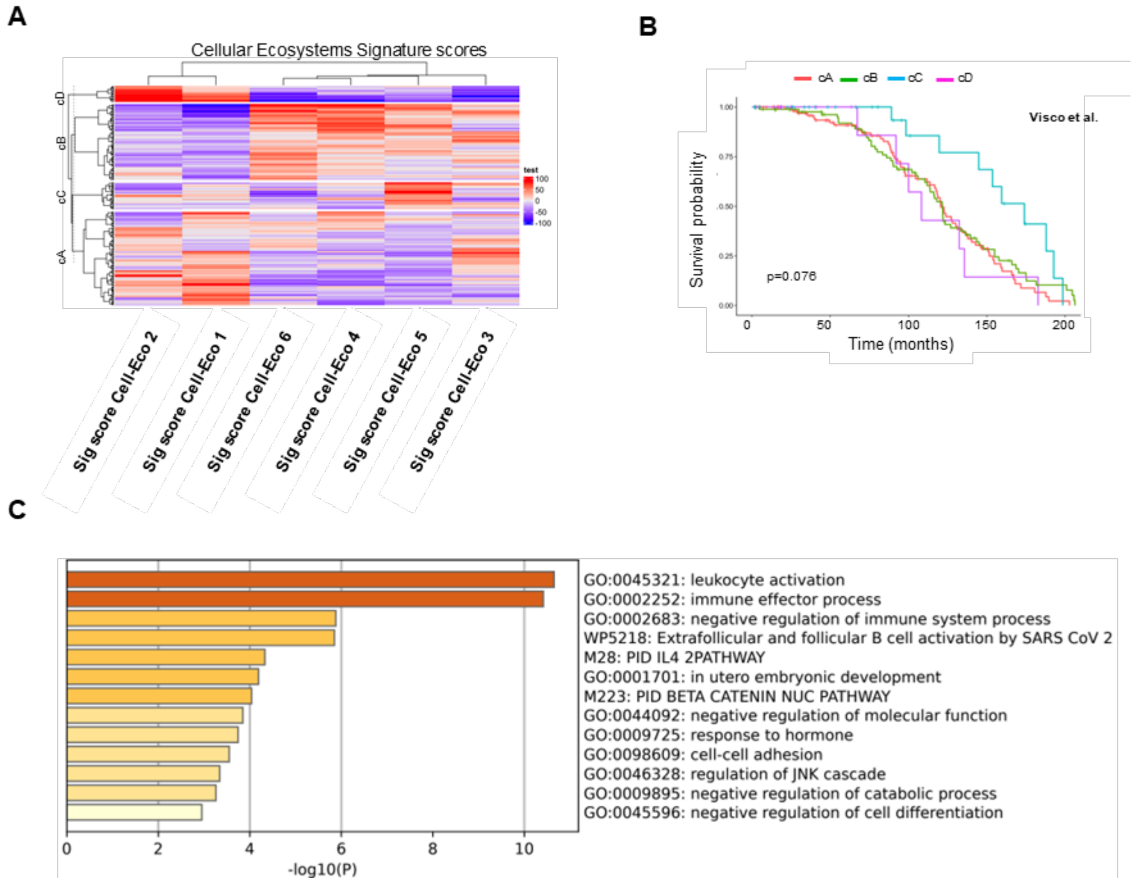


**Figure S4: Cell-Eco signature scores stratify patient survival in independent datasets. Related to Figure 3.**

A) Hierarchical clustering showing patients classification based on scaled Cellular Ecosystems signature scores in the Visco cohort of DLBCL RNAseq data (cA=187, cB=87, cC=31, cD=18). B) Kaplan-Meier curves with Long-rank test p-value showing overall survival of patients group stratified in Figure S4A. C) Functional pathway enrichment analysis of the top 30 genes with lower adjusted p-values , upregulated in Cell-Eco 5 compared to Cell-Eco 3. Only genes differentially expressed were considered (log2FC of 0.5 and $p < 0.05$)

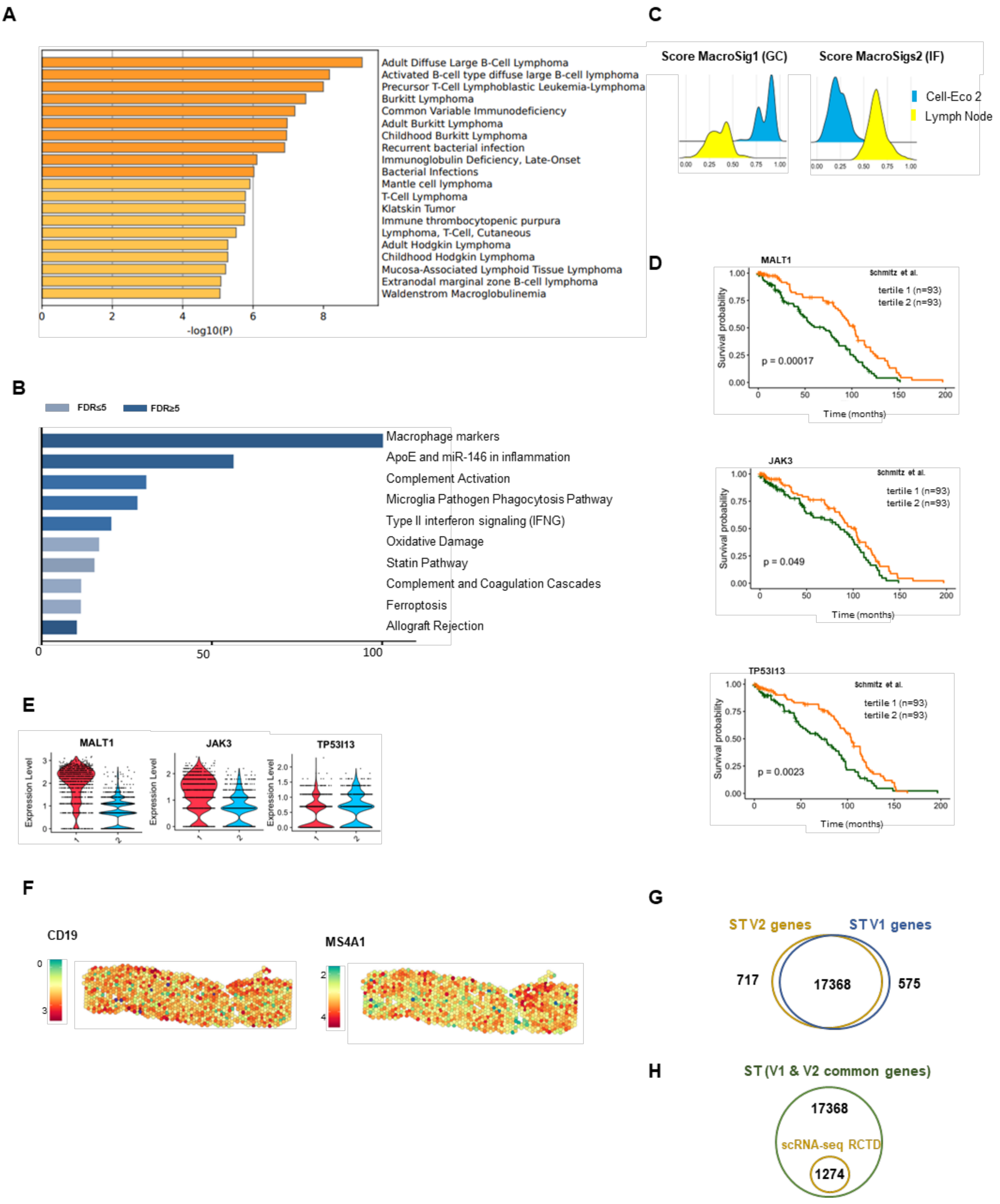
A
Adult Diffuse Large B-Cell Lymphoma
Activated B-cell type diffuse large B-cell lymphoma
Precursor T-Cell Lymphoblastic Leukemia-Lymphoma
Burkitt Lymphoma
Common Variable Immunodeficiency
Adult Burkitt Lymphoma
Childhood Burkitt Lymphoma
Recurrent bacterial infection
Immunoglobulin Deficiency, Late-Onset
Bacterial Infections
Mantle cell lymphoma
T-Cell Lymphoma
Klatskin Tumor
Immune thrombocytopenic purpura
Lymphoma, T-Cell, Cutaneous
Adult Hodgkin Lymphoma
Childhood Hodgkin Lymphoma
Mucosa-Associated Lymphoid Tissue Lymphoma
Extranodal marginal zone B-cell lymphoma
Waldenstrom Macroglobulinemia
-log10(P)
B
FDR≤5
FDR≥5
Macrophage markers
ApoE and miR-146 in inflammation
Complement Activation
Microglia Pathogen Phagocytosis Pathway
Type II interferon signaling (IFNG)
Oxidative Damage
Statin Pathway
Complement and Coagulation Cascades
Ferroptosis
Allograft Rejection
C
Score MacroSig1 (GC)
Score MacroSigs2 (IF)
Cell-Eco 2
Lymph Node
D
MALT1
Schmitz et al.
tertile 1 (n=93)
tertile 2 (n=93)
p = 0.00017
Survival probability
Time (months)
JAK3
Schmitz et al.
tertile 1 (n=93)
tertile 2 (n=93)
p = 0.049
Survival probability
Time (months)
TP53I13
Schmitz et al.
tertile 1 (n=93)
tertile 2 (n=93)
p = 0.0023
Survival probability
Time (months)
E
MALT1
JAK3
TP53I13
Expression Level
F
CD19
MS4A1
G
ST V2 genes
ST V1 genes
717
17368
575
H
ST (V1 & V2 common genes)
17368
scRNA-seq RCTD
1274

**Figure S5: Comparison of the transcriptomic profiles between Cell-Eco1 and Cell-Eco2 in ST DLBCL. Related to Figure 4.**

A) DisGeNET enrichment analysis of disease associated genes of the top 50 genes with lower adjusted p-values, upregulated in Cell-Eco 3 compared to Cell-Eco 5. Only genes differentially expressed were considered (log2FC of 0.5 and $p < 0.05$). B) Pathway enrichment of the top 50 genes with lower adjusted p-values , upregulated in Cell-Eco 5 compared to Cell-Eco 3. Only genes differentially expressed were considered (log2FC of 0.5 and $p < 0.05$). C) Module scores of external signatures indicating the location of macrophages in the lymph nodes between Cell-Eco2 and rLN spots with more than 10% of macrophages. D) Kaplan-Meier curves with Long-rank test p-value showing overall survival of patients divided by tertiles 1 and 2 for specific genes expression. E) Violin plots showing genes differentially upregulated in Cell-Eco 1 and Cell-Eco 2 and significant for survival analysis in two independent RNAseq DLBCL datasets. F) Spatial representation of gene expression organization of non-spatially autocorrelated genes on a representative ST DLBCL sample. G) Venn diagram of genes between Visium assays. ST data from the Visium V1 assay with direct placement (in blue) generated counts for 17,943 genes per sample, while the assay V2 or CytAssist-enabled assay generated counts for 18085 genes. 17368 genes are common to both assays. H) Venn diagram of genes common for the two assays (in green) and the genes from scRNA-seq data utilized by RCTD algorithm for cell-type mixtures deconvolution per spot (in yellow).
